# Common early-life exposure of the colorectal epithelium to the microbiome-derived mutagen colibactin

**DOI:** 10.64898/2026.11.16.766949

**Authors:** Henry N. Marshall, My H. Pham, Hyunchul Jung, Laura Humphreys, Yvette Hooks, Anh Phuong Le, Sarah Moody, Yichen Wang, Will Knight, Stephen Fitzgerald, Yan Shao, Soyeon Kim, Jiali Pang, Nicolas Wyvekens, Andrew Ramos, Calli Latimer, Rashesh Sanghvi, Ammal Abbasi, Claire Glemas, Kelly Spike, Komal Nayak, Rob Heuschkel, Ievgeniia Chicherova, Peter Campbell, Inigo Martincorena, Trevor D. Lawley, Ludmil B. Alexandrov, Tim H.H. Coorens, Raheleh Rahbari, Paul Brennan, Maria Dorn-Rasmussen, Vibeke Wewer, Johan Burisch, Matthias Zilbauer, Michael R. Stratton

## Abstract

Colibactin is a mutagen produced by bacterial strains in the colorectal microbiomes of some healthy people. It generates somatic mutations characterised by the mutational signatures SBS88 and ID18 in normal human colorectal epithelial cells *in vivo*. Increasing prevalence of colibactin mutagenesis during the 20^th^ century is hypothesised to have caused recent international increases in early-onset colorectal cancer incidence. Here, we establish the early-life prevalence, timing, mutation burdens and distribution through the colon of colibactin mutagenesis by sequencing normal colorectal epithelial cell genomes to detect SBS88 and ID18 in 90 healthy and inflammatory bowel disease affected children and adolescents. Colibactin-induced mutagenesis was present in ∼43%, with or without inflammatory bowel disease. It usually begins shortly after birth and ceases by age five years. Mutation burdens are higher in the left than right colon and show substantial variation, with some children displaying mutation burdens comparable to those of healthy non-exposed individuals greater than 60 years of age. The future public health implications of this common early life mutagen exposure will depend on clarification of its role in colorectal cancer causation.

## INTRODUCTION

Colibactin is a DNA damaging agent and mutagen which induces double strand DNA breaks, DNA crosslinking and adenosine alkylation in experimental systems^1–5^. It is produced by certain bacteria of the colorectal microbiome which are present in a proportion of healthy individuals, including *Escherichia coli* of the B2 phylogroup*, Klebsiella pneumoniae, Citrobacter koseri* and *Enterobacter aerogenes*, and which carry a 54-kb hybrid non-ribosomal peptide synthetase-polyketide synthase (*pks*) genomic cluster incorporating 19 genes^4,6,7,8^. Colibactin likely causes prophage induction and subsequent cell lysis of competing bacteria within the colorectal microbiome ecosystem^7,8^. Beyond its mutagenic capability, it may also induce cellular senescence and enhance tumour proliferation in human cells ^9–11^.

Co-culture of colibactin producing *pks+ E. coli* with human intestinal epithelial stem cell derived organoids, sequencing of organoid clones and matching of the mutation patterns generated indicated that colibactin is the causative agent of the single base substitution (SBS) and small insertion/deletion (ID) mutational signatures SBS88 and ID18 found in normal human colorectal epithelial cells *in vivo*^3,12,13,14^. SBS88 and ID18 are present in normal colorectal cells of some people but not others and are often generated during childhood^15^. They are enriched in early-onset compared to late-onset colorectal cancers, left colon/rectal cancers compared to right colon cancers and vary in prevalence internationally^12,16,17,18^.

The incidence of early-onset colorectal cancer has increased in many countries over the last four decades^19,20^. The cause is unknown but is suspected to be an exposure emerging during the course of the 20^th^ century which, at least in part, occurs during childhood and adolescence. The evidence that SBS88 and ID18 are caused by colibactin, are generated during the early decades of life, and are enriched in early-onset colorectal cancer has prompted the hypothesis that a global increase in the prevalence of early life colibactin exposure over the last century underlies the increase in early-onset colorectal cancer^18^. Evaluation of this hypothesis will require large-scale epidemiological studies of case-control and cohort designs. Here, we characterise in detail the features of colibactin mutagenesis in normal colorectal epithelial cells of children and adolescents as a foundation for these studies and to inform on the nature and scale of the future public health challenge it may represent.

## RESULTS

### Study participants and samples

90 children and adolescents (henceforth collectively referred to as “children”), aged 1.7-17.4 years (median 13.0), from the UK (n=73) and Denmark (n=17), of whom 54% were female, who had undergone colonoscopy in the period 2016-2024 as part of their clinical management, participated in the study (Fig. 1.B). Of 152 biopsies studied, 100 were from 57 children with normal colon mucosa (Supplementary Table 1) and 52 were from 33 children with Inflammatory Bowel Disease (IBD) (Fig. 1.B). Amongst children with normal colon mucosa, colonoscopy was predominantly indicated either to exclude IBD in the setting of gastrointestinal symptoms (n=43) or to investigate rectal bleeding (n=13). The majority of children with IBD (n=26) were treatment-naive at time of biopsy. Ascending and/or sigmoid colon biopsies were obtained from UK children (n=47 with both biopsies). A single transverse colon biopsy was obtained from 13 Danish children and four had additional biopsies of the terminal ileum, caecum, sigmoid colon, or rectum (totalling four biopsies per individual).

**Figure 1:**
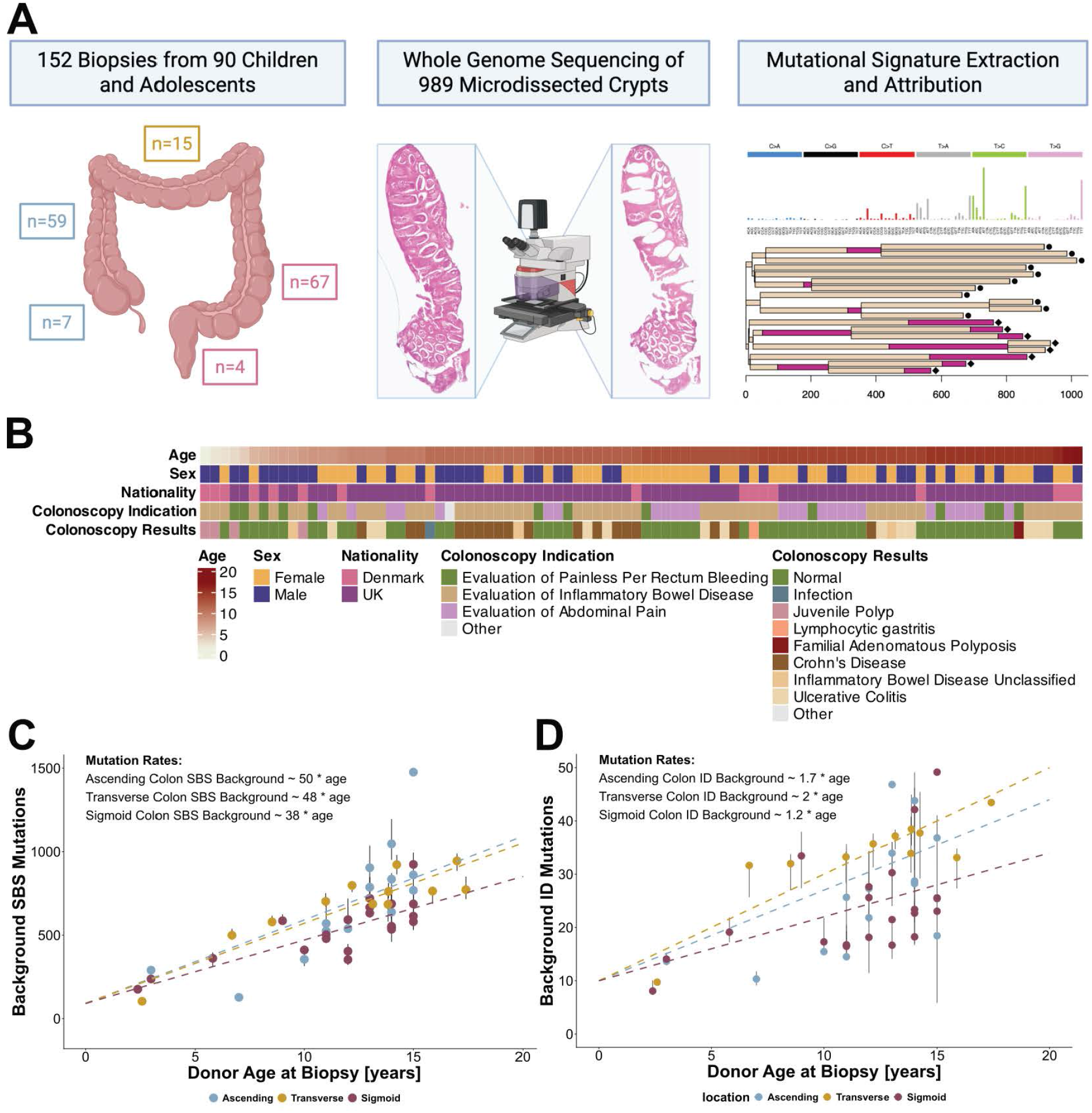
Study and Cohort Overview. ***A:** Study design. **B:** Heatmap of cohort overview for demographic and colonoscopy indication with results.* **C/D:** Background SBS and ID mutation burdens against age at time of biopsy. Background SBS mutations defined as SBS1, SBS5, and SBS18. Background ID mutations defined as non ID18 or IDA mutations. Median burden per biopsy denoted by dot with interquartile range (grey bar). Colour annotation denotes caecum/ascending colon (light blue), transverse colon (gold), and sigmoid/rectum colon (dark red). Dashed lines indicate estimates of mutation rates from linear mixed modelling.

### Mutational signatures

Crypts were microdissected from four caecum, 59 ascending colon, 15 transverse colon, 67 sigmoid colon, four rectum and three terminal ileum biopsies. Individual crypts (n=989) were whole genome sequenced to 20-fold median coverage and compared to DNA from an adjacent stromal tissue microdissection from the same child sequenced to median 33-fold coverage, yielding 780,455 somatic SBS and 49,024 ID in total^21,22^. SBS and ID mutational signatures were extracted from these mutation catalogues using both non-negative matrix factorisation (SigProfiler)^23^ and the hierarchical Dirichlet process (mSig HDP)^24^. A consensus set of extracted signatures was decomposed into reference mutational signatures from COSMIC v3.5 using SigProfiler Extractor. The mutation burdens of each signature in each crypt were estimated using MSA (Methods)^25^.

Five SBS mutational signatures were found: SBS1, SBS5, SBS18, SBS88 and SBS89. SBS1 is due to spontaneous deamination of 5-methylcytosine^26,27^, SBS18 is due to DNA damage by reactive oxygen species, and the mutational process underlying SBS5 is not well understood^28^. SBS1, SBS5 and SBS18 are endogenously generated mutational signatures, found in almost all crypts of all people, which accumulate throughout life in a linear correlation with age across multiple tissue types^15,29^. These have been combined in the following analyses as “SBSBackground”. SBS88 is due to colibactin^3,15^. SBS89 has been reported in normal colorectal epithelium and in colorectal cancers, is of unknown aetiology^18^ and was observed here in one individual.

Five ID mutational signatures were found: ID1, ID2, ID9, ID18, and IDA. ID1 and ID2 linearly accumulate with age^15,26,27^. ID9 is of unknown aetiology. ID18 is due to colibactin mutagenesis. IDA reflects the previously described ID counterpart to SBS89 and was characterised by 1bp deletions of T at non-homopolymer regions^15,30^. SBS89 / IDA was observed in one individual (PD67981; Fig 2.A). Mutations attributed to ID1, ID2 and ID9 have been combined and referred to as “IDBackground” mutations.

**Figure 2:**
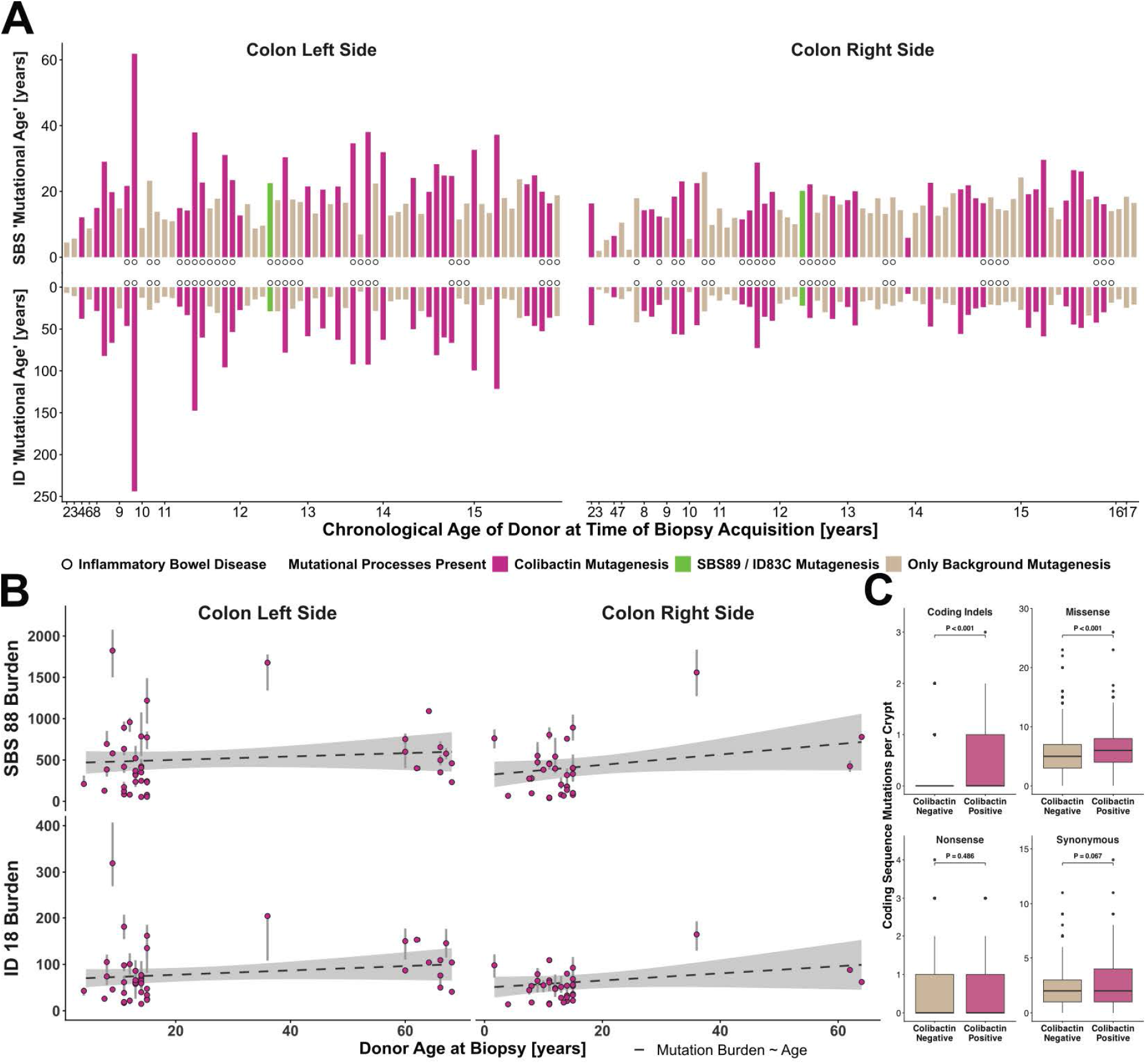
Prevalence and Mutation Burden of Colibactin Mutagenesis. **A:** Faceted bar plot showing the average SBS and ID ‘mutational age’ of each biopsy (y-axis) versus chronological age of the donor (x-axis). Biopsies are stratified into those of the left sided colon and those from the right sided colon. Bar plot indicates presence per donor of colibactin mutagenesis (pink), SBS89 / IDA mutagenesis (light green) or only background mutagenesis (light brown). **B**: Median SBS88 and ID18 mutation burden (light pink points) with interquartile range per biopsy (grey vertical bar) for colon left side and colon right side versus age at time of biopsy (x-axis) for children in present study and re-analysis of previously published data from adults^15^. Dashed line indicates estimate from linear mixed effects regression of mutation burden versus age, with mixed effects adjustment for biopsy and patient (95% confidence interval of regression denoted by grey shading). No association was observed between age at biopsy and SBS88 or ID18 mutation burden (SBS88 colon left side: p-value=0.400, SBS88 colon right side p-value=0.077, ID18 colon left side: p-value=0.187; ID18 colon right side: p-value=0.162). **C**: Box and whisker plot of number of indels, missense mutations, nonsense mutations, and synonymous for crypts that exhibited colibactin mutagenesis (colibactin positive; pink) and crypts that did not exhibit colibactin mutagenesis (colibactin negative; light brown). P-values derived from linear mixed modelling comparing colibactin positive and negative crypts adjusted for Bonferroni multiple testing correction.

Copy number changes and structural variants were rare. A copy number neutral loss of heterozygosity event affecting chromosome 17q was observed in a single crypt. A total of 63 unique structural variants were observed in 65 crypts, comprising 43 large deletions, eight tandem duplications, seven inversions and five translocations (Supplementary Table 4). There were no complex rearrangements or structural rearrangements involving known colorectal cancer genes. One individual (PD65118) demonstrated 10 unique structural variants. There was no enrichment of structural variants amongst children with colibactin mutagenesis in mixed effects regression accounting for age (0.06, 95% confidence interval (CI) [-0.02, 0.14] p-value=0.163).

### Background mutation rates

SBSBackground and IDBackground mutation rates were estimated using the mutation catalogues from the colon crypts of healthy children (ie without IBD or other colon-wide known disease) who did not exhibit SBS89/IDA or colibactin (SBS88/ID18) mutagenesis (n=33). Separate estimates for the ascending, transverse and sigmoid colon were obtained using linear mixed effects modelling. Rates of background mutagenesis were not estimated for the terminal ileum, caecum and rectum due to insufficient biopsies (n=2 for each site). Consistent with previous studies in adults^15,27^, linear associations were observed between age and burden of SBSBackground mutations for ascending (50 mutations/year 95% CI [38, 61]), transverse (48 mutations/year 95% CI [37, 59]) and sigmoid colon (38 mutations/year 95% CI [26, 50]) (Fig 1.C). Linear associations were also observed between age and background ID burdens for ascending (1.7 mutations/year 95% CI [1.1, 2.4]), transverse (2 mutations/year 95% CI [1.4, 2.7]) and sigmoid colon (1.2 mutations/year 95% CI [0.5, 1.8]) (Fig 1.D). The rates of background SBS and ID mutations in crypts from the transverse colon were comparable to crypts from the ascending colon (SBS p-value=0.315 ID: p-value 0.933), but higher than crypts from the sigmoid colon (SBS p-value=0.014 ID: p-value=0.027). Given the similarities of background mutation rates and the relatively small number of transverse colon biopsies, in subsequent analyses samples from the ascending and transverse colon have been combined as “right colon” and samples from the sigmoid colon termed “left colon”.

Adult-onset IBD is associated with elevated somatic mutation burdens^31^. Children with IBD also exhibited higher total SBS mutation burdens per crypt (mixed effects regression: 149 mutations 95% CI [59, 239] p-value=0.002; Supplementary Fig. 2) than children without, and mutational signature specific analyses showed that children with IBD had higher SBS1, SBS5, and SBS18 total mutation burdens per crypt (SBS1: 67 mutations 95% CI [108, 26], p-value=0.002; SBS5: 35 mutations 95% CI [68, 2], p-value=0.044; SBS18: 47 mutations 95% CI [73, 21], p-value=0.001)^31^. Children with IBD and without IBD demonstrated comparable total ID mutation burdens (3.9 mutations/year 95% CI [-4.9, 12.7] p-value=0.398; Supplementary Fig. 2)

### Prevalence and mutation burden of colibactin mutagenesis

The presence of colibactin mutagenesis in a given donor or biopsy was defined by the attribution of both SBS88 and ID18 mutations in at least one colorectal crypt. 39 children (43%) exhibited colibactin mutagenesis in at least one colon biopsy (67 biopsies were colibactin positive) (Fig. 2.A). Children from the UK had a higher prevalence of colibactin mutagenesis than children from Denmark (UK children: n=36 out of 73 [49%] versus Danish children: n=3 out of 17 [18%]; generalised linear model p-value = 0.031). The prevalence of colibactin mutagenesis was comparable amongst UK children with IBD (n=17 out of 33 [52%]) and without IBD (n=19 out of 40 [48%]; linear model p-value=0.686; minimum detectable difference at 80% power: 33 percentage points), as were the SBS88 and ID18 mutation burdens (SBS linear mixed effects p-value=0.395; ID linear mixed effects p-value=0.368) indicating that colibactin exposure does not contribute to the pathogenesis of IBD and that IBD does not influence colibactin mutagenesis.

Crypts from the left colon exhibited higher SBS88 and ID18 burdens than crypts from the right colon (SBS88 estimated difference: 76 mutations, 95% CI [26, 126] linear mixed effects p-value=0.003; ID18 estimated difference: 11 [3, 19] linear mixed effects p-value=0.004). Of the 27 colibactin positive children who had biopsies of both left and right colon, 23 showed colibactin mutagenesis in both biopsies, three in just the left colon, and one in just the right colon. Amongst colibactin positive biopsies, 59% (n=39 biopsies of 66) showed SBS88/ID18 in all crypts analysed, and the median number of crypts positive per biopsy was 5 ([QR [3, 8]] median of 100% of crypts positive). In only three donors was an individual positive in just one crypt. Within biopsies, the standard deviation in mutation burden across crypts was typically about 25% of the mean for SBS88 and 32% of the mean for ID18 (cohort median coefficient of variation SBS88: 0.25 IQR [0.16, 0.33] and for ID18: 0.31 IQR [0.23, 0.39]). Analysis of biopsies from the caecum, ascending colon, transverse, sigmoid and/or rectum from two Danish children indicated the presence of colibactin mutagenesis in all colonic sites (Fig. 3.A/3.B) but absent in two of the three terminal ileum crypts in PD69095 (Fig. 3.A).

**Figure 3:**
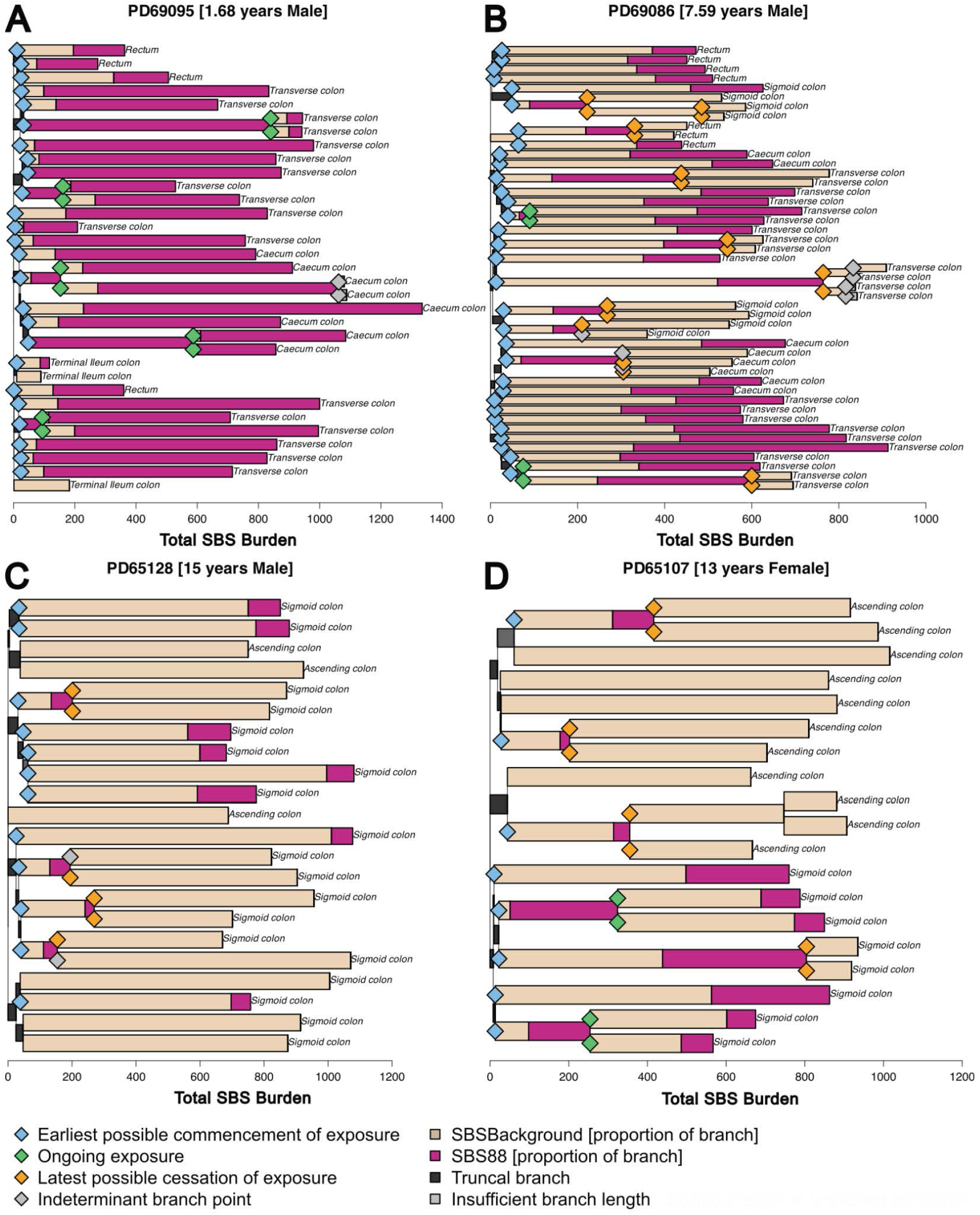
Phylogenetic Analyses of Colibactin Mutagenesis. **A/B/C/D:** Example of four phylogenetic trees. Each branch is denoted by the proportion of background SBS mutations (light brown) and proportion of SBS88 mutations (pink). The start of the first positive branch marks the earliest possible commencement of exposure for a crypt (blue diamond). Blue diamonds are only depicted if a branch began after time point 0 for informative branches. Timepoints of ongoing exposure are represented by bifurcations at which both the branch preceding and the branches following the bifurcation exhibit SBS88 (green diamonds). Timepoints for latest possible cessation of exposure are indicated by bifurcations where the preceding branch exhibits SBS88 and the subsequent branch is negative for SBS88 on pruned and unpruned bootstrapped signature weights and exceeds 44 mutations. Indeterminate branch points are bifurcations where the preceding branch exhibits SBS88 but the subsequent branch was negative for SBS88 but exhibited SBS88 unpruned attribution or was less than 44 mutations long. The x-axis for each tree depicts the cumulative number of SBS mutations in the tree.

SBS88 and ID18 mutation burdens varied substantially between colibactin-exposed children (SBS88: 28 - 2311 mutations; ID18: 4 - 489 mutations). To convey the impact of colibactin mutagenesis, the “mutational age” of each crypt was estimated, i.e. the chronological age at which the observed number of mutations would be expected if just background mutational signatures had been operative. The presence of colibactin mutagenesis increased the SBS mutational age of crypts in both the left colon (mixed effects estimate: 11 years, 95% CI [7, 15] p-value<0.001) and the right colon (mixed effects estimate: 4 years, 95% CI [2, 6]; p-value=0.001) (Fig. 2.A). The effect of colibactin mutagenesis on mutational age was greater when considering ID mutations for both the left colon (median difference: 49 years, 95% CI [34, 64]) and the right colon (median difference 18 years, 95% CI [13, 23]) (both mixed effects p-values<0.001). Certain children showed particularly high levels of colibactin mutagenesis. For example, PD69095, biopsied at age 1.7 years, had an SBS mutational age of 16 years and ID mutational age of 45 years and PD68018, biopsied at age nine years, had an SBS mutational age of 62 years and ID mutational age of 244 years.

With respect to the consequences for encoded proteins, colibactin positive crypts exhibited more missense (p-value<0.001) and coding sequence ID (usually translation frameshifting) mutations (p-value<0.001) than colibactin negative crypts (Fig. 2.C). However, the numbers of nonsense mutations were similar due to the preponderance of T>C substitutions in SBS88 which do not cause translation termination codons (Supplementary Fig. 4). No likely driver mutations in cancer genes were found^16^.

Inherited mutations in POLE and POLD1 (which encode replicative DNA polymerases) and MUTYH (which encodes an element of the repair machinery for DNA damage caused by reactive oxygen species) cause elevated mutation rates in normal colorectal epithelial cells^32,33^. To contextualise the effect of colibactin mutagenesis, the mutation burdens observed in the current study were compared to those expected from the somatic mutation rates reported in previous analyses of individuals with inherited POLE, POLD1, MUTYH mutations^34,35^. Most children with colibactin mutagenesis (67%) exhibited colon epithelium mutation burdens similar to, or greater than, those estimated for children with inherited causes of accelerated mutagenesis of the same age (Supplementary Fig. 5). Although this analysis highlights the implications of colibactin exposure on somatic mutagenesis, conclusions regarding cancer predisposition are beyond the scope of this analysis.

### Timing of colibactin mutagenesis

The prevalence of colibactin mutagenesis in children was comparable to that in previously reported adults (43% of children versus 37% of adults; generalised linear model p-value=0.622; minimum detectable difference: 20 percentage points). Both SBS88 and ID18 burdens displayed no associations with age across the range of children and adults studied (SBS88 colon left side: 2.0 mutations/year 95% CI [-2.8, 6.8] p-value=0.400; SBS88 colon right side: 6.3 mutations/year 95% CI [-0.7, 13.2] p-value=0.077; ID18 colon left side: 0.5 mutations/year 95% CI [-0.2, 1.2] p-value=0.187; ID18 colon right side: 0.8 mutations/year 95% CI [-0.3, 1.8] p-value=0.162; Fig. 2.B)^15^. The consistency of prevalence and mutation burdens across age ranges studied, and its presence amongst children under age 10, is consistent with mutagenesis starting and usually stopping during childhood.

To refine estimates of the timing of initiation and cessation of colibactin mutagenesis a phylogenetic tree of colorectal crypts for each individual was constructed and mutational signatures attributed separately to each branch. Starting from the origin of the tree and proceeding through to the tips, the first branch positive for SBS88 and ID18 represents the time window during which colibactin mutagenesis commenced, with the earliest possible timepoint being the start of the branch (blue diamonds; Fig. 3), and the latest possible timepoint for commencement of exposure being the termination of the branch. Timepoints at which colibactin mutagenesis is ongoing (assuming that colibactin mutagenesis is continuous) are represented by bifurcations at which both the branch preceding and the branches following the bifurcation exhibit SBS88 and ID18 (green diamonds; Fig. 3). Time windows for cessation of colibactin mutagenesis are indicated by branches which exhibit SBS88 and ID18 and for which the subsequent branches are negative for SBS88 on pruned and unpruned bootstrapped signature weights and exceeded at least 44 mutations long (orange diamonds; Fig. 3; Supplementary Fig. 7), with the start of the SBS88 and ID18 positive branch indicating the earliest timepoint at which colibactin mutagenesis ceased and the end of the branch indicating the latest timepoint. The chronological age of each branchpoint was estimated from the cumulative total number of mutations attributed to background mutational signatures (SBS1, SBS5, SBS18) prior to the branchpoint.

Most phylogenetic trees included one or more short initial branches representing fewer than 50 mutations. Mutational signature attributions to such small datasets are not reliable. To address this problem composite spectra of these short early branches from all trees were constructed. These showed only the background mutational signatures SBS1 and SBS5, without evidence of SBS88, even in children with colibactin mutagenesis in later branches (Supplementary Fig. 6). These early branches likely represent mutations arising *in utero* and are consistent with initiation of colibactin mutagenesis after birth when the colorectal microbiome is acquired.

Colibactin mutagenesis usually starts early in postnatal life. Informative branchpoints from the phylogenetic trees of 32 children showed that the median age after which mutagenesis could have started was 0.5 years (IQR [0.3, 0.9]; Fig. 4 blue diamonds) and in no instance did the first colibactin positive branch for a crypt commence after the age of two. Analysis of the end points of the first positive branches indicated that the median age by which colibactin mutagenesis must have started was 2.3 years (IQR [1.3, 4.2]; Fig. 4 orange/green diamonds). The timepoints for the earliest and latest possible ages of commencements were enriched for earlier in life than expected under a permutation-based null distribution accounting for tree topology (Kolmogorov-Smirnov p-value < 0.001; Supplementary Fig. 9). The earliest single timepoint by which colibactin mutagenesis must have started in a crypt was 0.4 years in PD69095. All colibactin positive children with informative branchpoints by age five showed colibactin mutagenesis by age five (Fig. 4). Consistent with these observations, adults with informative branches (n=5) demonstrated a median latest age of commencement of 7.3 years (range [4.6, 7.6]), although more refined estimates in this group were precluded by insufficient branchpoints (Supplementary Fig. 10). Collectively, the results are consistent with colibactin mutagenesis commencing during the first few months to early years of life.

**Figure 4:**
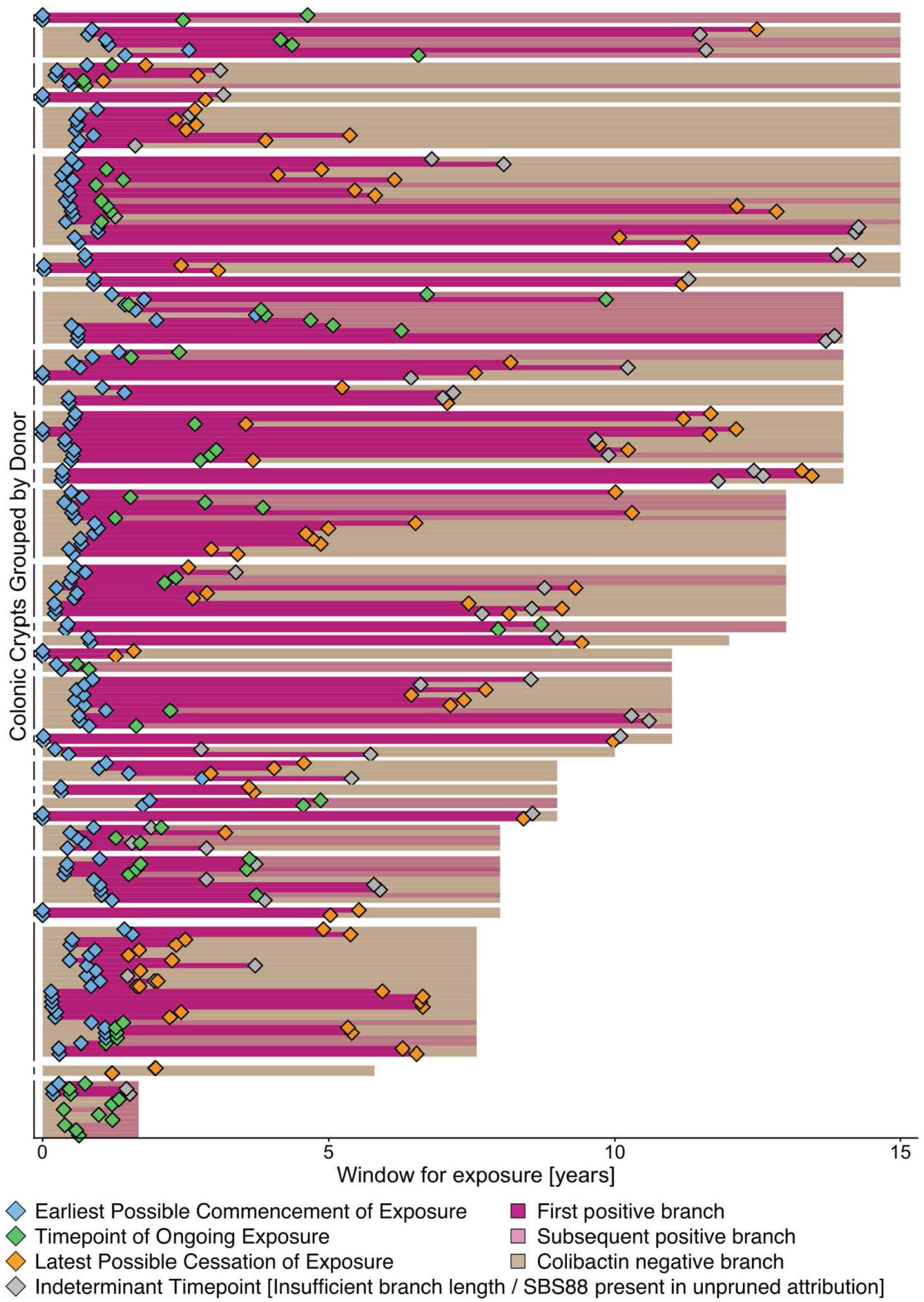
Timing of Colibactin Mutagenesis. Swimmer plots depicting estimated informative time points (diamonds) of exposure to colibactin mutagenesis for 32 children (y-axis) with informative phylogenetic trees. The length of the bar indicates the age at which the biopsy was acquired. Dark pink indicates the first positive branch in a crypt and lighter pink indicates second positive branch in a crypt phylogeny. Blue diamonds denote the earliest possible age of exposure commencement for a given crypt, green diamonds denote ages of definitively ongoing exposure within a crypt (indicated by colibactin mutagenesis present both before and after a phylogenetic bifurcation), orange diamonds denote the latest possible time point of exposure cessation for a given crypt, and grey diamonds indicate ‘indeterminant timepoints’ whereby no SBS88 was attributed on the subsequent branch using pruned MSA results, but either the branch was deemed of insufficient length or SBS88 was present in the unpruned attribution.

Colibactin mutagenesis frequently ceases during childhood. Amongst crypts which demonstrated cessation of mutagenesis, the median age by which cessation must have occurred was 5.1 years (IQR [2.9, 7.8]; Fig. 4.B yellow diamonds). The earliest age by which colibactin mutagenesis must have ceased in a crypt was estimated at 1.1 years in PD65120 (Fig. 4). The latest age for continuing was 9.8 years (PD65109). The ratio of branchpoints showing ceased compared to continuing colibactin mutagenesis was 10:38 (0.26) before age 2, 35:23 (1.5) between 2-5 years, 30:7 (4.3) between 5-9 years and 13:0 at 10-15 years (mixed effects regression for association between timepoint status and age p-value < 0.001). Similarly, two adults demonstrated cessation by age 5, and all adults showing cessation demonstrated cessation in at least one crypt by age 10 (cohort median earliest timepoint: 6.1 years [range 0.8, 9.8]; Supplementary Fig. 10). Thus, colibactin mutagenesis usually ceases before age five years and almost always before age 10.

The duration of colibactin mutagenesis may vary between children. PD69086 had five crypts that all showed continuation between the estimated ages of 1.1 to 1.5 years, and then cessation in 20 crypts with a median age of 3.7 (IQR [2.2, 6.0]; Fig. 3.B). By contrast, PD65109 showed three branchpoints indicating ongoing colibactin mutagenesis between ages 4 and 7, and one branchpoint at age 13 that showed cessation. The duration of colibactin mutagenesis may also vary between anatomical locations. Whereas no crypts from the right colon demonstrated continuation beyond three years of age (n=37 timepoints), the median age of continuation for left sided crypts was 3.8 years (IQR [2.2, 4.8], n=37 timepoints; Fig. 4 green diamonds). PD65107 and PD65128 illustrated this difference. PD65107 showed cessation in six right colon crypts before age 5, but continuation in a left colon crypt at age 4.3 and then cessation in two left colon crypts at age 10.5 (Fig. 3.D). In PD65128, colibactin mutagenesis was entirely absent from right sided crypts, and estimated to have ceased before age six years in six crypts of the left colon (four crypts showed cessation by age four) (Fig. 3.C). Crypts which showed a timepoint of continuation exhibited a higher SBS88 mutation burden than crypts that did not (Wilcox Rank p-value<0.001).

## DISCUSSION

Over the last decade colibactin has been revealed as a microbiome-derived mutagen to which a significant proportion of the global human population is exposed^18^. The consequences of colibactin exposure are uncertain but a contribution to colorectal cancer risk, and in particular early-onset colorectal cancer, has been hypothesised^3,15,18^. Previous work indicates that colibactin mutagenesis occurs in childhood^15^. This study establishes at high resolution the features of colibactin mutagenesis in children, healthy and with IBD, to support future examination of this hypothesis.

Participants in the study had undergone colon/rectal biopsy. This procedure in children was prompted only by significant symptomatology, which raises the possibility of cohort ascertainment bias influencing the results and generalisation of the conclusions. Some children suffered from IBD, which increased background mutation rates, but did not show association with the prevalence or burden of colibactin mutagenesis. Most children did not have microscopic evidence of colon mucosal abnormality and, in many, the diagnosed cause of symptoms (for example, a single juvenile polyp) was unlikely to engender consequences for the whole colon mucosa. Samples in such cases were from unaffected colon. While significant ascertainment bias appears unlikely, however, it cannot be completely excluded.

SBS88 and ID18 mutation burdens due to colibactin exposure were higher in the normal left colon than right colon, consistent with their higher prevalence in left colon/rectum cancers^18^. The reason for this distribution is not known but plausibly is related to differences in population size of *pks*^+^ organisms influencing the amplitude of colibactin exposure in the different locations. It is notable that the increased early-onset colorectal cancer incidence over the last three decades is also enriched for left colon/rectal cancers.

Colibactin mutagenesis starts early in life at estimated timepoints consistent with the immediate postnatal period during which the newborn’s colorectum is colonised by the microbiome. Its presence amongst individuals as young as 1.7 years of age and five individuals younger than age 10 indicated that this exposure must commence during the early years of life. Phylogenetic analysis illustrated that mutagenesis proceeds in an apparently continuous fashion before ceasing, usually before age five years and almost always before age 10. Restarting of colibactin mutagenesis later in life is not excluded by the results reported here but has not been observed in other studies. The mechanism underlying cessation of colibactin mutagenesis is unclear. It could be due to disappearance of the *pks*+ microbe from the colorectal microbiome. However, studies of adult microbiomes report the presence of *pks*+ bacteria later in life^9,36,37^. Alternatively, changes in the intracolonic environment leading to a reduction in the production of colibactin by *pks*+ bacteria throughout childhood^7^, or establishment of physical barriers, such as mucus, that prevent the adhesion of colibactin to the epithelial cell necessary for genotoxic activity^38^ may lead to reduction or cessation in mutagenesis. Stool samples from participants in this study were not available to evaluate these hypotheses further. The analysis of non-human reads revealed high levels of contamination accompanied by low levels of gut related bacterial taxa, which is likely due to artefacts and contamination associated with DNA extraction, the necessity for 10-12 rounds of PCR amplification for this method of low input whole genome sequencing, and library preparation^22,39–41^. Similarly, maternal or neonatal medical history was unavailable, precluding the analysis of epidemiological determinants of exposure. Future studies of SBS88 and ID18 mutation burdens in colorectal epithelial cells of children together with detection of *pks*+ organisms in their microbiomes and detailed clinical histories may clarify the mechanism of cessation.

The mutation burdens conferred by colibactin mutagenesis are highly variable between children. The factors underlying this variation are not understood but could plausibly be related to variation in size of the population of *pks*+ bacteria in the colorectal microbiome between children. Alternatively, *pks*+ gene transcription is known to be influenced by factors such as intestinal inflammation, or levels of glucose, iron and inulin^42–44^. Danish individuals had a lower prevalence of colibactin mutagenesis, which may reflect geographic variation in exposure or cohort ascertainment biases. The burden and prevalence of colibactin mutagenesis were comparable between children with histologically normal biopsies and children with IBD, indicating that these are independent processes.

Some children with colibactin mutagenesis exhibit markedly elevated mutational ages of their colon epithelial cells compared to their chronological ages. In particular, the high ID mutational ages observed reflect the propensity of colibactin mutagenesis to generate a proportionally higher ID compared to SBS mutation rate than background mutational signatures. This characteristic may be important in its potential contribution to colorectal carcinogenesis. Somatically acquired, biallelic *APC* gene mutations, which inactivate the APC protein through premature truncation, are common in colorectal cancer and are considered to be the first step of neoplastic transformation from normal crypt to cancer^45,46^. It is plausible that early life exposure to colibactin accelerates the process of transformation by increasing the probability of acquisition of the first inactivating *APC* driver mutation. SBS88 is, however, relatively ineffective in generating premature translational termination codons due to the preponderance of T>C mutations in the signature (Supplementary Fig. 4). By contrast, almost all ID18 mutations will engender translational frameshifts and consequently premature termination codons. Analysis of colorectal cancers suggests that ID18 is a common cause of *APC* ID mutations and confirms that SBS88 makes a limited contribution to translation termination codons but contributes to the cluster of *APC* splice variants at c.835-8A>G^18,47,48^. Therefore, the high ID mutation rate of colibactin mutagenesis may primarily be responsible for a putative contribution to colorectal carcinogenesis.

The prevalence of colibactin mutagenesis in children born in the UK and Denmark between 2001 and 2022 was ∼43%. Although there was substantial variation in the mutation burdens conferred, >10% of all children showed higher ID mutation burdens in normal left colon cells as a result of colibactin mutagenesis than observed in 50-year-olds without colibactin mutagenesis. In such children, the childhood ID mutation burdens are similar to those estimated in children with elevated colorectal cell mutation rates due to rare inherited mutations in *POLE*, *POLD1* and *MUTYH* which are known to confer elevated risks of colorectal cancer. Although this does not signify that a child with colibactin induced mutagenesis would specifically have the colorectal cancer risk of a 50 year old or individual with *POLE* mutation, it highlights the magnitude of this additional mutation burden in early childhood. Overall, the nature of the mutations constituting the SBS88 and ID18 signatures, their prevalence, their timing, their burdens, and the proportions of colorectal cells affected will together determine their capability to induce *APC* mutagenesis and confer an elevated colorectal cancer risk for children who are exposed.

The hypothesis that colibactin induced mutations contribute to colorectal cancer risk requires formal epidemiological evaluation through studies comparing the burdens of SBS88 and ID18 mutations in normal colorectal epithelial cells from colorectal cancer cases with those in matched healthy controls. Similarly, it was beyond the scope and power of this current study to evaluate whether the prevalence of colibactin mutagenesis has increased amongst young people. Such an analysis would require sequencing of normal colorectal epithelium or colorectal cancer samples from large numbers of individuals exposed during the course of the last century. The results reported here establish the foundation for this next generation of studies by illustrating that colibactin exposure is common amongst children, commences and ceases within the first years of life, and often confers substantial mutation loads.

## METHODS

### Study cohort

The prevalence, burden and timing of colibactin mutagenesis were assessed in a combined cohort of 152 colonic epithelial biopsies from 90 children from the United Kingdom and Denmark. One hundred and twenty-three biopsies of the ascending or sigmoid colon from 73 of these children were obtained through collaboration with the Cambridge University Hospitals. Study approval was granted by the East of England-Cambridge South Research Ethics Committee (reference number: 17/EE/0265). All patients underwent colonoscopy as part of their routine clinical diagnostic work-up for the investigation of gastrointestinal symptoms. Forty children had both macroscopically and histologically normal colonic mucosa. The indication for colonoscopy amongst these children was for the exclusion of IBD in the assessment of abdominal pain (n=26) or painless rectal bleeding (n=13). The clinical indication for one child was unavailable on review. The majority (n=38 children and adolescents) had a complete resolution of symptoms and can therefore be classed as ‘healthy controls’. Although a small number of patients had other related pathology (e.g. juvenile polyp n=1 child, familial adenomatous polyposis n=1 child) biopsies taken from those patients were obtained from a region of macroscopically and histologically normal colon. Thirty-three children and adolescents demonstrated histopathological changes indicative of IBD (Supplementary Table 1). Age, sex and medical history were recorded for all participants. A further 29 biopsies from 17 children were obtained through collaboration with the Copenhagen IBD Inception Cohort Study (3 terminal ileum, 4 caecum colon, 15 transverse colon, 3 sigmoid colon and 4 rectum biopsies)^49^. The Copenhagen IBD Inception Cohort Study protocol was approved by the Ethics Committee of the Capital Region of Denmark (H-20065831) and the Danish Data Protection Agency (P-2020-1065). These samples were specifically. The indication for colonoscopy in these Danish children was for the evaluation of IBD. These children all had macroscopically and histologically normal biopsies (Supplementary Table 1). Of the children with histologically normal biopsies in this cohort, one later developed Crohn’s disease. Sample biopsies were obtained from all participants at the time of colonoscopy.

Written informed consent for all samples in this study was obtained from all donors or their legal guardian in accordance with the Declaration of Helsinki. No statistics were performed to predetermine cohort size.

### Tissue sample preparation

Colonic samples from both cohorts were snap-frozen at the time of colonoscopy. Colonic biopsy samples were fixed with PAXgene fixative (PreAnalytiX) and paraffin-embedded to preserve tissue morphology^22^. Samples were sectioned at 16µm onto 4mm PEN membrane slides (Leica) with subsequent Gill’s Haematoxylin and Eosin staining. Individual crypts were isolated with laser-capture microdissection (LMD7000, Leica) and collected in separate wells of a 96-well plate. The minimum microdissected tissue for each crypt was 10,000µm^2^. For each colonic biopsy, a section of ∼100,000µm^2^ of adjacent stromal or submucosal tissue was dissected for use as a matched normal. Cell lysis of crypt biopsies was performed using the Arcturus PicoPure Kit (Applied Biosystems) as per the manufacturer’s instructions and incubated at 65°C for 12 hours^22^. Each well was subjected to low-input DNA library preparation^22^, and samples with DNA concentration > 5ng/µL underwent whole genome sequencing. Whole genome sequencing of colonic crypt samples was performed using the Illumina NovaSeq X platform, using 150bp paired-end sequencing reads targeting a coverage of ∼30x. The median deduplicated coverage of crypts was overall lower than the matched normal representing inherited sequence variation (median 20x versus 33x). This difference reflected the difference in amounts of DNA obtained (median concentration 15ng/uL for individual crypts versus 65ng/uL for the normal controls) that arose due to less cellular material being available from each crypt due to their spatial constraints compared to the normal tissue control. Sequences were aligned using BWA-MEM (version 0.7.17) and mapped to the GRCh38 human reference genome^50^. Assessment of contamination and concordance with matched normal was conducted using Conpair^51,52^. Assessment of evenness of coverage was performed by calculating a mean over median score per crypt. Crypts were excluded if contamination level was >3%, concordance with matched normal was < 98%^51,52^, or coverage < 10x^52^.

### Single base substitution variant calling and filtering

Single base substitution variants were called using a validated consensus-based calling approach using the CaVEMan algorithm^53^ (version 1.15.2) and Strelka2^54^ (version 2.9.10)^48,55^. CaVEMan copy number options were set to major copy number 5 and minor copy number 2. Both CaVEMan and Strelka2 were run using adjacent stromal tissue as a matched reference normal. A series of post-processing filters were then applied to remove potential artefacts arising from either mapping errors or PCR-associated cruciform artefacts^22,56^. First, a read-aware artefactual variant flagging tool, *Hairpin2*^57^, was applied to identify 1) variants which appear with anomalously regular positional distribution in supporting reads, 2) reads with poor signal-to-noise ratio as per alignment score, 3) PCR-associated sequencing errors^29^. This was run using default parameters^57^. CaVEMan variants were then intersected with variants also called by Strelka2 and only those variants called by both algorithms were kept. A pileup of the mutant and wild-type reads for all mutations across all samples from a given individual was constructed. Only those variants with a VAF >0.05, depth >7 and at least 4 mutant reads, mapping quality of 30 and base quality of 30 in at least one sample from a given donor were kept. Finally, a beta-binomial model was applied to identify artefacts that may be present at low but consistent frequencies across samples^58^. Following filtering, variant allele frequency histograms were reviewed and crypts with a median VAF < 0.3 were excluded due to polyclonality. The full pipeline used for variant filtering is publicly available at: https://github.com/Phuong-Le/lcmb.

### Indel variant calling and filtering

Short insertion and deletion (ID) variants were called using a similar pipeline to single base substitutions. Variants were called using both Pindel (v3.11.0) and Strelka2 (v2.9.10) and only variants called by both were kept^54,59^. *Hairpin2* was again run, with modified parameters to account for analysis of ID variants. A pileup of variants was once again run to apply a beta-binomial filter excluding low frequency artefactual variants.

### Copy number and structural variant calling

Copy number profiles were generated using ASCAT NGS (v4.5.0) and manually reviewed. Samples demonstrating marked aberrancy, often due to low coverage, were excluded. A pileup of copy number segments across all donors was constructed to assess for any chromosomal arm copy number changes. Structural variants (SVs) were called with GRIDSS^60^. Specifically, GRIDSS (version 2.9.4) was used with default settings. SVs larger than 1kb in size with QUAL ≥250 were included. For SVs smaller than 30kb, only SVs with QUAL ≥300 were included. Furthermore, SVs that had assemblies from both sides of the breakpoint were considered only if they were supported by at least four discordant and two split reads. SVs with imprecise breakends (i.e., the distance between the start and end positions > 10bp) were filtered out. To remove low-input-WGS-specific artefacts (i.e., calls supported only by PCR-duplicate-like reads), we further filtered out SVs for which the standard deviation of the alignment positions at either end of the discordant read pairs was smaller than five. To capture potential germline SVs and artefacts, SVs found in the matched normal or in at least three different donors in the in-house panel of normals were removed.

### Mutational signature extraction and attribution

Mutational signatures were extracted and decomposed using a consensus approach for both SBS and ID mutations. For both SBS and Indels, mutational signatures were extracted by creating a mutation signature matrix of unique variants per patient^58,61^. Extraction of mutational signatures were performed using non-negative matrix factorisation methodology, SigProfiler^23^, and a Bayesian Hierarchical Dirichlet Processes, mSigHDP^23,24^. SigProfiler was run using default parameters and maximum 10 signatures. mSigHDP was run using 5000 burnin, K.guess=10. The average stabilities and mean cosine distance for all proposed solutions from SigProfiler were reviewed. The optimum solution from SigProfiler contained two signatures (SBS96A and SBS96B). SBS96A decomposed into: SBS1, SBS5, SBS18, and SBSS96B decomposed into SBS1, SBS5, SBS88. mSigHDP extracted four high confidence signatures, referred to as HDP1, HDP2, HDP3, HDP4 (Supplementary Fig. 1). SBS96A and HDP2 shared a cosine similarity of 1, and SBS96B and HDP1 shared a cosine similarity of 0.98. HDP3 had a cosine similarity of 0.28 with SBS96A and 0.74 with SBS96B. HDP3 was present only in samples that harboured HDP1. HDP 4 showed low cosine similarity with either SBS96A or SBS96B (0.3 and 0.19 respectively), It arose predominantly in PD67981 where it contributed to 34% of unique mutations (3439 out of 10,083). It decomposed into SBS89, and significantly improved the spectra fitting for PD69781 when included (expectation maximisation with 1000 bootstraps p-value<0.001)^62^. Signature decomposition for both the outputs from SigProfiler and msigHDP was performed using the Analyze.decompose_fit() from SigProfilerAssignment (v0.1.4). (https://github.com/hmarshall14/ChildhoodColibactinExposure).

ID signatures were similarly extracted by a pileup of unique mutations for each donor using the 83-channel SigProfiler classification based on insertion/deletion, degree of homology, and length of mononucleotide tract. Three de novo mutational signatures were extracted using mSigHDP (ID83A, ID83B, ID83C). ID83A decomposed into ID1, ID2, ID9 (cosine similarity: 0.99). Given that ID1 and ID2 accumulate with age, this likely reflected background mutagenesis. ID83B reflected ID18. ID83C did not decompose into any COSMIC (v3.5) signatures. It was characterised by 1bp deletions of T at non-homopolymer runs and decomposed into the previously described counterpart to SBS89 (cosine similarity of 0.99 compared to IDB from Lee-Six et *al.*)^15,30^. For the purposes of this analysis, ID83C was referred to as IDA

SBS and ID mutation signatures were attributed to each crypt using MSA with 2000 simulated samples^25^. MSA performs mutational signature assignment on a single-sample basis using Non-Negative Least Squares with optimization using simulations and bootstrapping^25^. Bootstrapped ‘pruned’ results from MSA outputs were used for downstream analysis. Pruning refers to the process of ‘zero-ing’ the estimated weight of a signature in a sample if the lower bound of the 95% confidence interval for the estimated weight of the signature crosses zero following bootstrapping of simulated data. We elected to use MSA following a review of multiple signature attribution modalities: SigProfiler, MuSiCal, MSA, mmSig, and sigfit (Supplementary Fig. 3). Following MSA attribution, the median per crypt cosine similarity for SBS attribution was: 0.97 IQR [0.93, 0.98] (correlation median: 0.97 IQR [0.96, 0.98]), and for ID attribution was: 0.94 IQR [0.90, 0.96] (correlation median: 0.93 IQR [0.89, 0.96]).

### Characterising prevalence of colibactin mutagenesis

The presence of colibactin mutagenesis in either a biopsy or donor was defined by the attribution of both SBS88 and ID18 to at least one crypt. The requirement of both SBS88 and ID18 was implemented to mitigate the risk of over estimation, particularly during ID attribution (Supplementary Fig. 3). Only three individuals demonstrated SBS88 and ID18 in a single crypt. If a crypt exhibited either SBS88 or ID18 but not the other, it was deemed colibactin negative. Comparisons of the prevalence of colibactin mutagenesis between IBD and normal children, and between UK and Danish children were performed using generalised linear modelling accounting for age, sex and the presence of IBD (in UK/Danish comparisons). Comparisons of prevalence between children and adults were performed using generalised linear modelling accounting for sex and IBD status.

### Mutation burden analysis

All SBS and ID mutation burdens were adjusted for coverage and variant allele frequency, by calculating a sensitivity score that estimates the likelihood of detecting an SBS variant with at least four mutant reads (or five reads for ID variants)^56,63^. This score is calculated by running 100,000 simulations derived from a set of Bernoulli tests with a success probability equal to the median VAF of the sample^56,63^. The number of tests is derived from a Poisson distribution with lambda equal to the mean coverage.

Linear mixed effects regressions estimated the age-associated rate of background SBS and ID mutagenesis from those histologically normal children who did not exhibit colibactin mutagenesis, familial adenomatous polyposis, or SBS89/IDA (n=33 children). Background SBS rates were estimated using the combined attribution of SBS1, SBS5, and SBS18, which have known age associated correlations^15,63^. Background ID rates were estimated using the combined burden of non-ID18 or IDA mutations in these crypts, thereby approximating to the total ID burden in crypts from these children. Background rates were estimated for ascending, transverse and sigmoid colon crypts but not terminal ileum, caecum or rectum due to insufficient biopsies for these sites. A structured stepwise approach was applied to estimate the age associations of background SBS mutations for the ascending, transverse and sigmoid colon. First a base model with age as a fixed effect and a random intercept per donor was fitted to the SBS background mutations per crypt (*SBS Background mutations ∼ age + (1 | patient)*). An interaction-only term for crypt biopsy location was then included to determine the per location estimated effect of age on background SBS mutations (*SBS Background mutations ∼ age:location + (1 | patient)).* This model outperformed the base model (ANOVA p-value<0.001). The same approach was used for estimates of age associations for background ID mutations. All mixed effects modelling was performed using the *lmerTest* package.

We next assessed the influence of IBD amongst colibactin negative cases. A linear mixed effects model incorporating an additional fixed effect for IBD status, ‘*IBD_status,’* was fitted to background SBS mutations (model: *SBS background mutations ∼ age:location + IBD_status + (1 | patient)).* This model outperformed a base model which included the interaction term of age and location but did not account for IBD status (ANOVA p-value<0.001). Secondary analyses for SBS signature specific associations with IBD were fitted using the same model structure. The influence of IBD on ID mutation rates was similarly assessed using the model *ID background mutations ∼ age:location + IBD_status + (1|patient).* A final model tested whether the presence of colibactin mutagenesis per crypt influenced the burden of background SBS mutations. The inclusion of a binary value ‘colibactin_status’ (positive / negative) per crypt did not improve the model performance (ANOVA p-value=0.205). Because the majority of IBD individuals in this study were undergoing diagnostic colonoscopy and the date of onset of symptoms was unknown, no analysis of the effect of duration of symptoms was performed^31^.

The SBS88 and ID18 mutation burdens of children’s colons were compared to adult colonic crypts from previously published data^15^. Consensus calling for SBS and ID variants was performed using either CaVEMan and strelka2 or Pindel and strelka2. Variants were subject to the same filtering pipeline as the children’s data (https://github.com/Phuong-Le/lcmb). The previously identified set of signatures from this cohort (SBS1, SBS2, SBS5, SBS13, SBS18, SBS88, SBS89, ‘SBSC’, and ‘SBSD’) were attributed using MSA.^15^ Comparison of the prevalence of colibactin mutagenesis between children and adults was performed using generalised linear modelling. Association between age and SBS88 or ID18 mutation burden was tested using a linear mixed effects model: mutation burden ∼ age:location + (1|patient). Terminal ileum and rectum biopsies were excluded from this analysis because they were absent from the adult population. Mutational ages of colonic crypts were estimated by dividing the total number of mutations present in a crypt against the left colon or right colon (as appropriate for each crypt) specific estimates of the rate of background mutagenesis per year derived from colonic crypts from healthy individuals without colibactin mutagenesis.

### Phylogenetic tree construction

Phylogenetic trees of colonic crypts for each individual were constructed using the Sequoia framework^58,64^, in short by applying a maximum parsimony algorithm to the depth count and variant count matrices of the final set of filtered single base substitutions for each individual (MPBoot v1.1.0). Single base substitutions were mapped to the children’s tree branches using a maximum likelihood approach (treemut R package https://github.com/nangalialab/treemut) and visualised using *ggtree* (v3.14.0) and *ape* (v5.8-1). Mutational signatures were attributed to each branch of the phylogenetic trees using MSA. Branch lengths were adjusted for each branch’s sensitivity score for detecting a mutation with 4 supporting reads. Sensitivity scores for tips of the tree were given as the sensitivity score for that crypt, whereas sensitivity scores for internal branches *‘A’* with two child branches ‘*B*’ and ‘*C’* was given as *(1 – Sensitivity_A) = (1 - Sensitivity_B) x (1 – Sensitivity_C)*. Estimates of the timing of breakpoints were determined by dividing the number of background SBS mutations up to a point by the estimated rate of background mutagenesis for a given crypt. Ninety-five per cent confidence intervals on age of breakpoint were estimated by using an exact Poisson test on the cumulative background mutations up to the breakpoint and the total background mutations. A branch was deemed colibactin positive if SBS88 was attributed in the pruned MSA attribution. The earliest possible commencement was defined as the start of the first positive branch in the evolution of a crypt, and the latest possible commencement was defined as the termination of the first positive branch. Timepoints of continuation of exposure were branchpoints where both the preceding and subsequent branches were colibactin positive. Timepoints of cessation were defined as branchpoints where the preceding branch was colibactin positive but the subsequent branch was colibactin negative. These subsequent branches were referred to as ‘post-cessation branches.’

We applied a stringent criterion to define post-cessation branches. These branches were required to demonstrate an absence of SBS88 in the unpruned bootstrapped attribution, and be of sufficient length to ensure that absence of attribution was not an artefact of short branch length. We derived a threshold from a simulated dataset. Five thousand five hundred samples with a burden of 5 to 630 mutations were simulated, of which 500 had no SBS88. Proportions of SBS1, SBS5, SBS18 in simulated data were sampled from the 5th to 95th percentile of SBS1/SBS5/SBS18 proportions observed in SBS88-negative paediatric crypts. Attribution using MSA was performed on simulated data (Supplementary Fig. 7.A), and the probability of detecting SBS88 with 1% and 5% false-negative rate was modelled using logistic regression (Supplementary Fig. 7.B). The minimum branch length threshold was estimated as the intersection between the curve for the 5% false-negative rate and the 5th percentile for branch length in our observed data. This intersection corresponded to 44 mutations (Supplementary Fig. 7.C). Timepoints at which the ‘post-cessation branch’ either showed SBS88 on unpruned attribution or had < 44 mutations were deemed ‘indeterminant timepoints.’ Several secondary analyses supported the absence of colibactin mutagenesis amongst these post-cessation branches. Several secondary analyses validated the absence of colibactin mutagenesis amongst these post cessation branches. The analyses illustrated that post cessation branches had comparable rates of indel mutations per year to branches from colibactin negative crypts (P=0.137), no enrichment of any trinucleotide contexts compared to branches from colibactin negative crypts, and clustered on principal components analysis with branches from colibactin negative crypts (Supplementary Fig. 8).

A permutation-based analysis evaluated whether the early ages of commencement of exposure observed in this study were simply an artefact of sampling young individuals. Branches in phylogenetic trees were randomly assigned as SBS88 positive or negative. SBS1, SBS5, and SBS18 were unchanged between permutations to facilitate timing analysis. 10,000 permutations were performed and the observed values were compared to the permuted values. A Kolmogorov-Smirnov test assessed whether the observed distribution was less than the permuted distribution. Mixed effects regression assessed the relationship between age and enrichment of continuation timepoints (relative to cessation timepoints) to points early in life. The following model was fitted to the timepoints of continuation or cessation: *timepoint ∼ age_at_timepoint + age_at_biopsy + (1|patient).* Age at biopsy was included to account for bias of younger individuals in the study. Random intercept per patient accounted for clustering of multiple timepoints per individual.

### Mutational driver assessment

A pileup of unique mutations per donor was constructed and coding mutations were annotated using the dNdScv package^65^. Rates of coding indels, missense, nonsense and synonymous mutations between colibactin positive and negative crypts were compared by fitting a generalised linear model with mixed effects for colibactin mutagenesis status and age, and a random effect of patient and biopsy location using the *lmerTest* package. Identification of cancer driver mutations was undertaken by running Cancer Genome Interpreter^66^.

### Microbial analysis of colibactin presence

Whole genome sequencing BAM files were converted to FASTQ format. Low-quality reads were removed using fastp^67–69^. Microbial read processing and analysis from human WGS data followed a best-practice workflow as recently described by Salzberg and colleagues^70^. Depletion of host genome was performed by excluding reads that mapped to either: GRCh38, T2T-CHM13.v2.0, or the 47 Human Pangenome Reference Consortium dataset^67,68^. To detect the presence of *Escherichia coli,* the primary colibactin-carrying species, remaining non-human reads were first classified against a curated, comprehensive microbial database (Microbial2023; available at https://benlangmead.github.io/aws-indexes/k2) using KrakenUniq^71^, a *k*-mer-based metagenomic classifier optimised for detecting microbial signals in WGS datasets with low microbial biomass^70,72^. In parallel, non-human reads were also mapped to the CHOCOPhlAnSGB database using MetaPhlAn4^73^, a widely used metagenomic classifier that leverages clade-specific marker genes for taxonomic classification.

Evaluation of the association between colibactin mutagenesis and microbiome was performed by assessment of the relative abundance of bacterial taxa from consensus read counts from KrakenUniq and MetaPhlAn4, and assessment of the presence of the colibactin-encoding pks gene cluster. Bacterial taxa were annotated as likely contaminants, gut taxa, or unknown. Contaminants were taxa that had previously been documented as reagent, laboratory or environmental contaminants^40,41^. Within these taxa, we flagged a subset that were biologically impossible within colonic epithelium (i.e. those present in extremely halophilic archaea, soil- or marine-restricted clades). Gut taxa were those documented in the Unified Human Gastrointestinal Genome collection^74^. Remaining taxa were marked as unknown.

Colonic samples demonstrated high levels of contaminant reads and low coverage of gut taxa. 76% of all classified non-human reads were assigned to contaminant taxa and only 5% to gut taxa (Supplementary Figure 10). The three most abundant taxa were all contaminants (combined assignment: 40.1% of reads). Furthermore, the relative abundance of contaminant-associated reads exceeded 0.5 in 95% of crypts and all crypts displayed biologically impossible gut taxa, indicating that this was a cohort-wide phenomenon. Only 179 crypts demonstrated sufficient coverage to confidently call at least one gut bacterial taxon (defined by mapped coverage greater than 0.05X, median taxa called for those 179 samples: [IQR: 1, 1; max: 42]). Taken together the results indicate that excessive contaminants and low coverage of gut taxa preclude evaluating differences in microbiome composition between colibactin mutagenesis positive and negative samples.

## ACKNOWLEDGEMENTS

This work was supported by the Wellcome Trust grants 206194 and 220540/Z/20/A and by a European Research Council Synergy Grant ECCE (grant agreement #101224262). H.M. is supported by a John Monash Scholarship. J.B. is supported by the EU Horizon Europe Program grant miGut-Health: personalised blueprint of intestinal health (101095470) and Novo Nordisk Clinical Emerging Investigator Grant NNF20OC0059377. We would like to thank L. O’Neill, A. Vasisht, S. Austin-Guest and the staff of the Wellcome Sanger Institute Sample Logistics, Genotyping, Pulldown, Sequencing and Informatics facilities for their support with sample management, laboratory work and analytical pipelines. We would like to thank P. Scott for his advice in laser capture microdissection. We thank the donors and their families for their participation in this study.

## DISCLAIMER

Where authors are identified as personnel of the International Agency for Research on Cancer/World Health Organization, the authors alone are responsible for the views expressed in this article and they do not necessarily represent the decisions, policy or views of the International Agency for Research on Cancer /World Health Organization. J.B. reports grants from AbbVie, Janssen-Cilag, MSD, Takeda, Tillots Pharma, Bristol Myers Squibb, and Novo Nordisk Foundation; personal fees from AbbVie, Janssen-Cilag, Celgene, MSD, Pfizer, Takeda, Tillots Pharma, Bristol Myers Squibb, Samsung Bioepis, Pharmacosmos, Ferring, Galapagos, Eli Lilly, Dr Falk Pharma, Celltrion, Zealand Pharma and Orion Pharma. L.B.A is a co-founder, CSO, scientific advisory member, and consultant for Acurion (formerly io9), has equity and receives income. The terms of this arrangement have been reviewed and approved by the University of California, San Diego in accordance with its conflict of interest policies. LBA is a compensated member of the scientific advisory board of Inocras. L.B.A’s spouse is an employee of Hologic, Inc. L.B.A. declares U.S. provisional applications with serial numbers: 63/289,601; 63/269,033; 63/366,392; 63/412,835; 63/966,993 as well as international patent application PCT/US2023/010679. L.B.A. is also an inventor of a US Patent 10,776,718 for source identification by non-negative matrix factorization. L.B.A. further declares a European patent application with application number EP25305077.7. I.M., P.C. and M.R.S. are academic co-founders, consultants and equity holders of Quotient Therapeutics.

## DATA AVAILABILITY

Whole genome sequencing data has been deposited on the European Genome-Phenome Archive (EGA) with accession code EGAD00001015830. Processed data are available in the Supplementary Tables. Code and additional files used to construct analyses can be found on GitHub (https://github.com/hmarshall14/ChildhoodColibactinExposure).

## SUPPLEMENTARY FIGURE

**Supplementary Figure 1:**
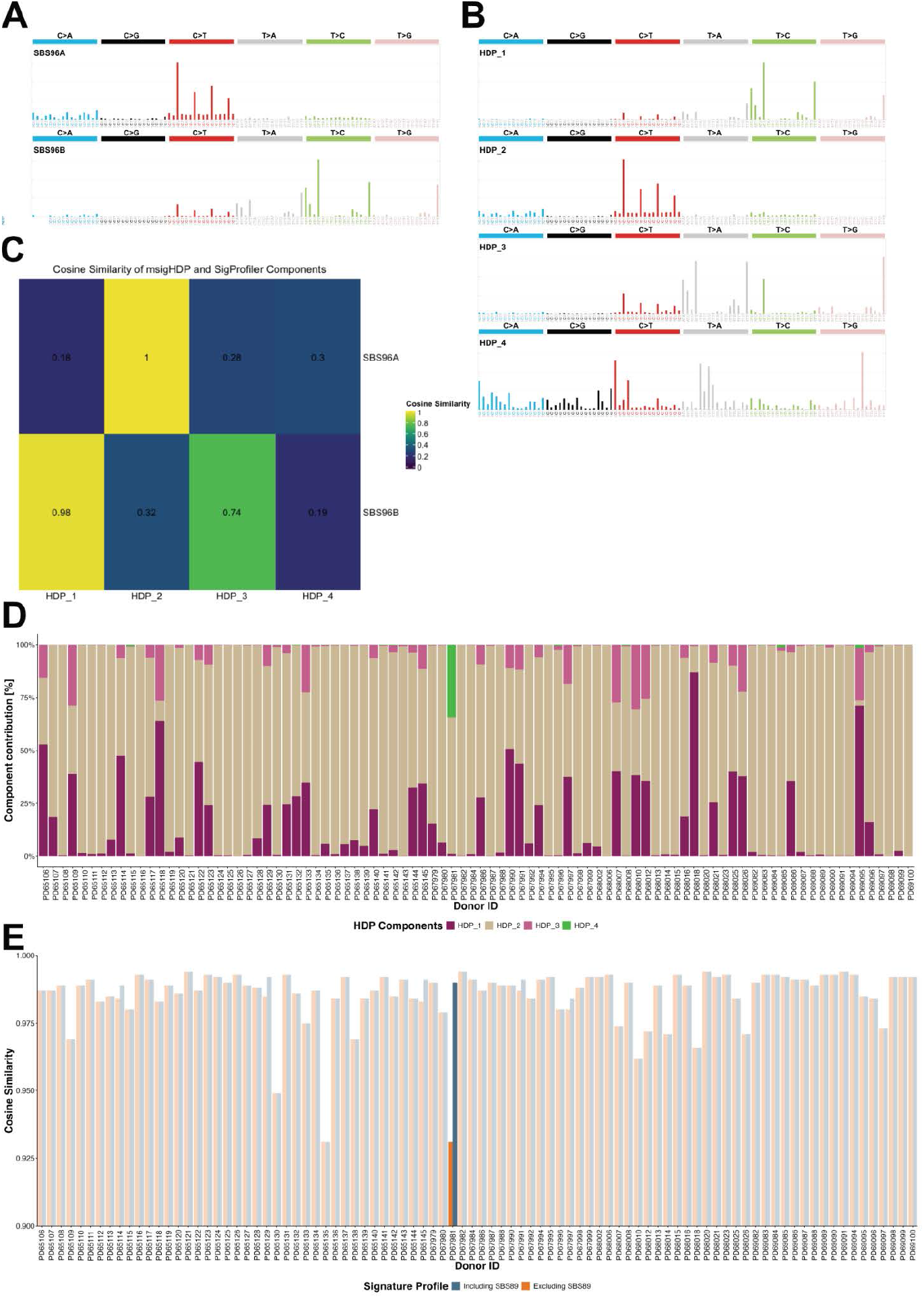
Mutational signature extraction. **A:** Signatures extracted using SigProfiler. **B:** Signatures extracted using msigHDP. **C:** Cosine similarity of signatures extracted by SigProfiler (SBS96A, SBS96B), versus signatures extracted by msigHDP (HDP_1,HDP_2, HDP_3, HDP_4). **D:** Proportion of mutations attributed to each extracted signature in each donor, depicting 34% of mutations in PD67981 attributing to HDP_4. **E:** Comparison of cosine similarity with (blue) and without (orange) the inclusion of SBS89 into the signature fitting profile. Including SBS89 into the attribution for PD67981 significantly improved its fit (expectation maximisation p-value<0.001).

**Supplementary Figure 2:**
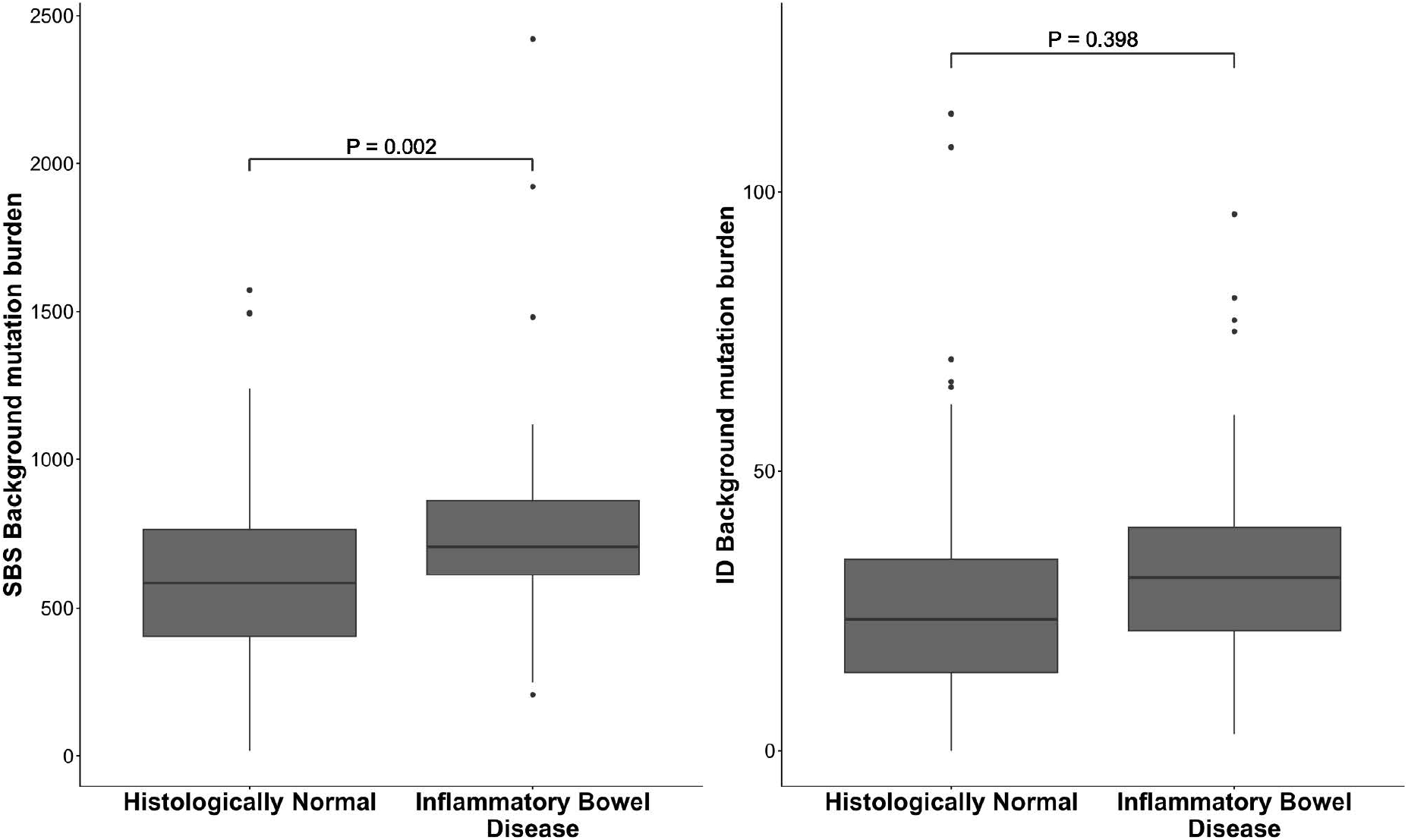
Comparison of Background Mutation Burden for Inflammatory Bowel Disease and Histologically Normal Children. Boxplot and whisker plot showing the distribution of background SBS and ID mutations per crypt for inflammatory bowel disease and histologically normal children and adolescents. P-value derived from linear mixed effects accounting for age and location of crypt biopsy.

**Supplementary Figure 3:**
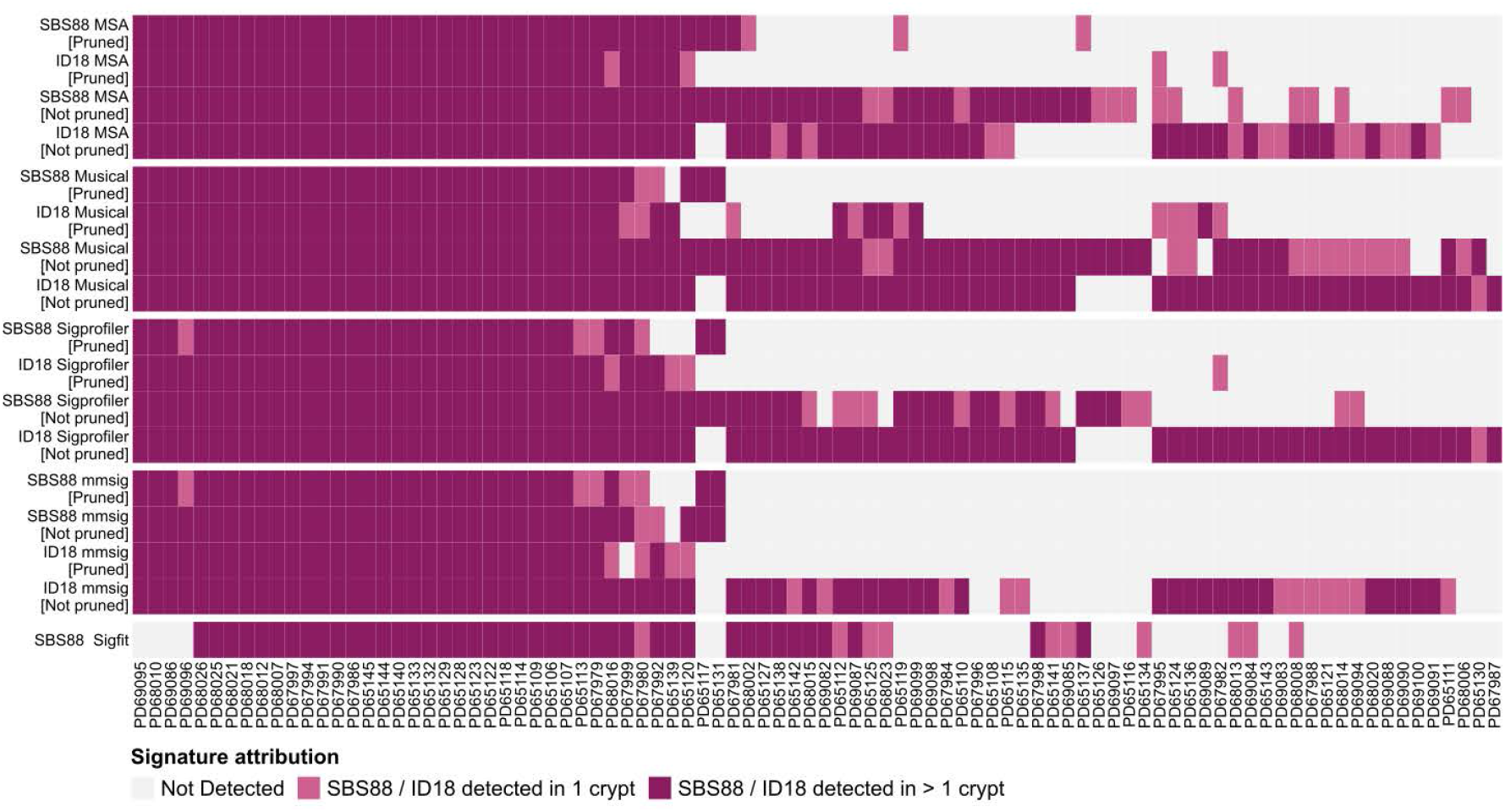
Comparison of detection of colibactin mutagenesis across different signature fitting algorithms. Heatmap depicting the presence of SBS88 or ID18 in either no, one (light pink) or two or more crypts (dark pink) in each of the 90 children in this study for different methods of signature attribution (MSA, Musical, SigProfiler, mmSig, and Sigfit) with and without pruning of bootstrapping results. “Pruning” refers to the process whereby the estimate for the signature weight is zero-ed if the 95% confidence interval crosses 0.

**Supplementary Figure 4:**
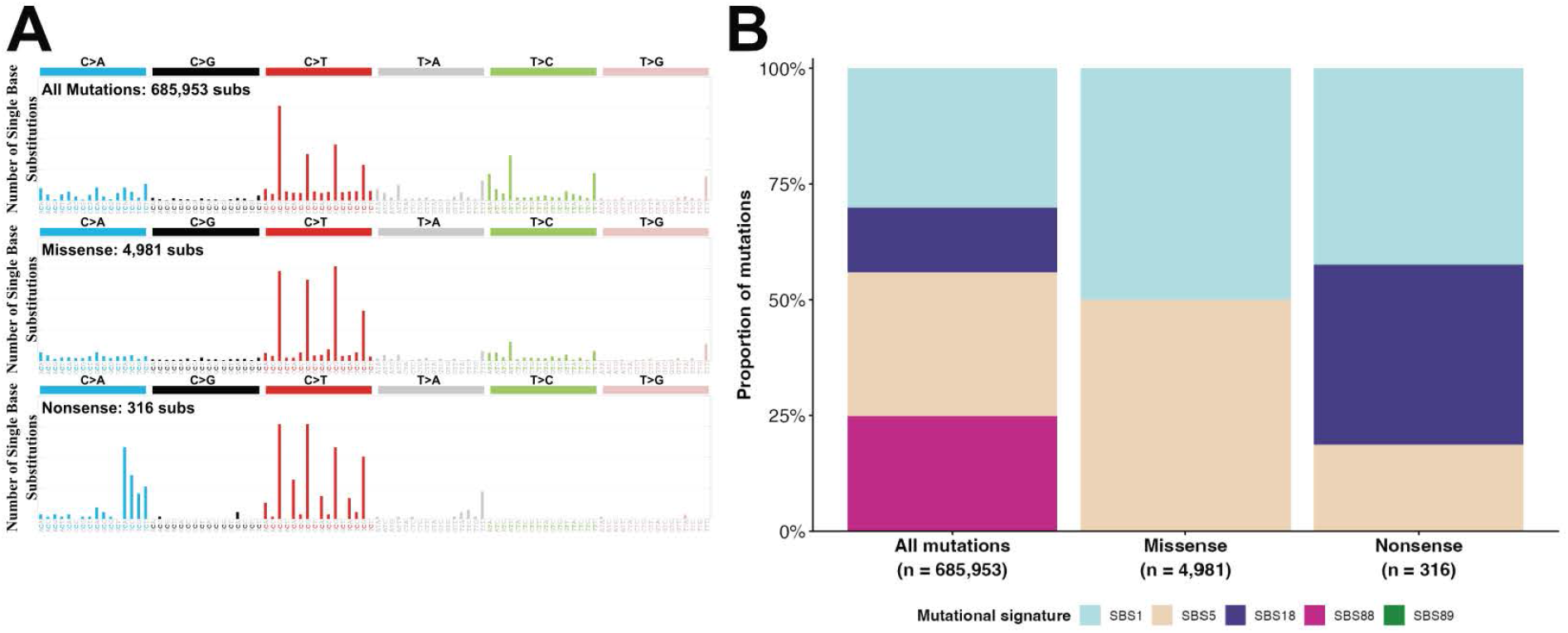
Probability of coding mutations from SBS88. **A:** Mutational spectra pileup of all unique mutations (top), all unique missense mutations (middle) and all unique nonsense mutations (bottom) among children’s colonic crypts. **B:** Bar plot depicting signature attribution to: all unique mutations, all unique missense mutations and all unique nonsense mutations. No nonsense or missense mutations were attributed to SBS88. Mutations were annotated using *dNdS* package. Attribution was performed using *SigProfiler* Assignment.

**Supplementary Figure 5:**
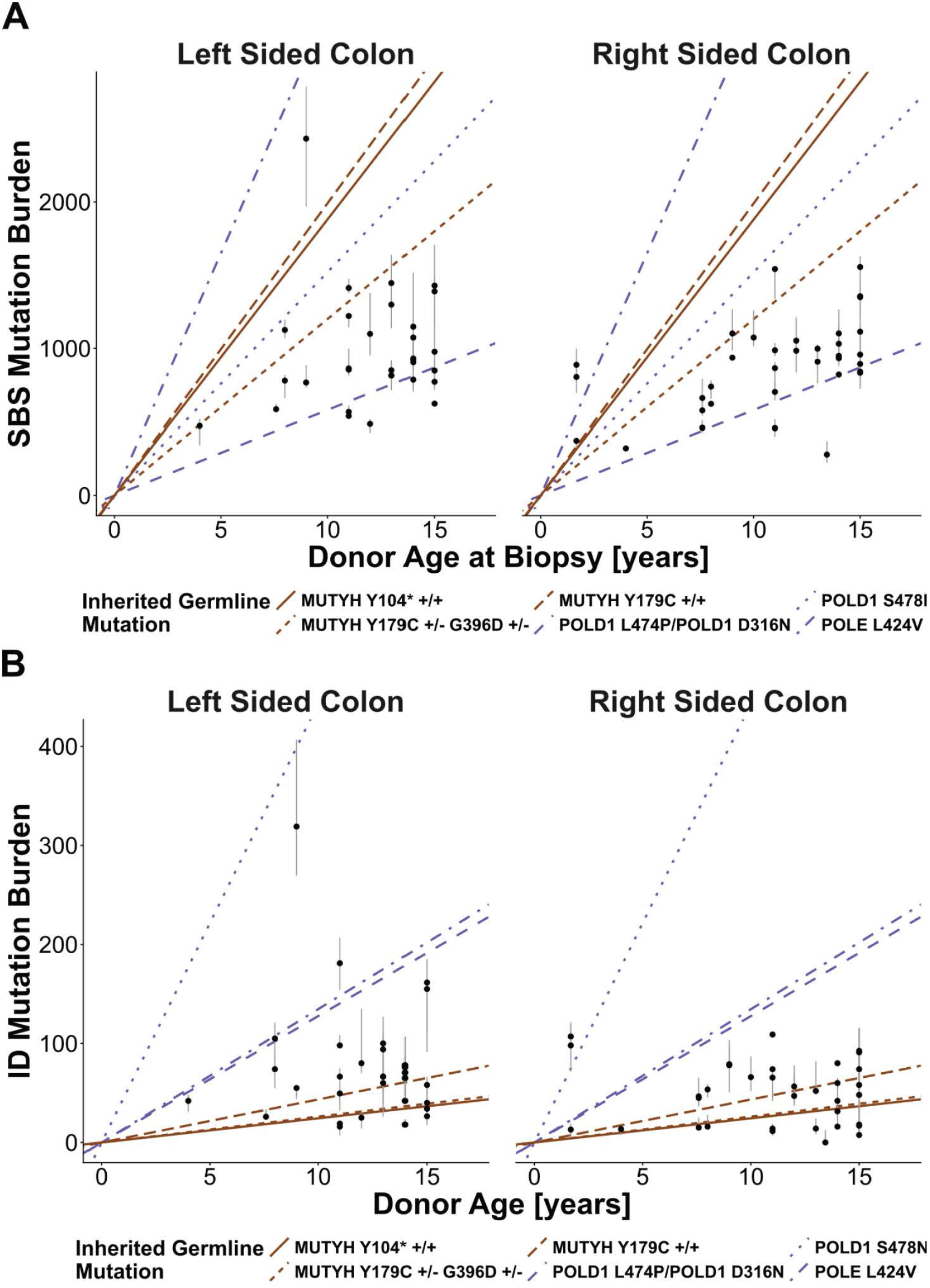
Mutation Burdens for Colibactin Positive Individuals Relative to Rates of Mutagenesis in Inherited Cancer Predisposition Syndromes. **A/B**: Total SBS (A) and ID (B) mutation burden for colibactin positive biopsies (black dot: median; grey line: interquartile range) from left sided and right sided colon. Previously estimated rates of SBS and ID mutation burdens for children with inherited mutations in *MUTYH* (brown lines)^34^ and *POLE* and *POLD1* (purple lines) genes^35^.

**Supplementary Figure 6:**
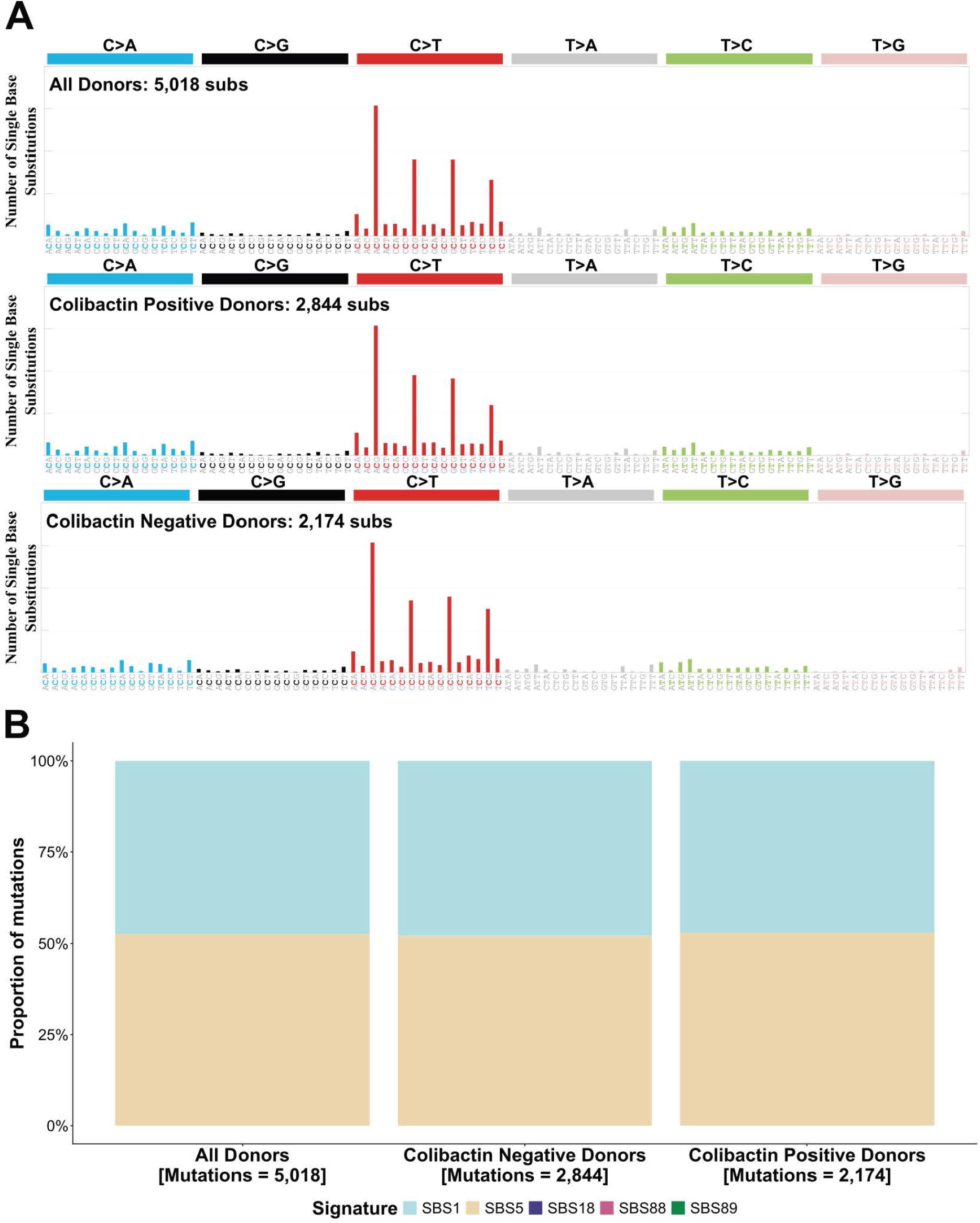
Mutational Signature Extraction from Early Phylogenetic Branches. **A:** Mutation spectra for early phylogenetic branches from all children (top), for early phylogenetic branches from children without colibactin (middle) and for early phylogenetic branches from children with colibactin mutagenesis (bottom) **B:** Attribution of mutational signatures extracted from study cohort (SBS1, SBS5, SBS18, SBS88 and SBS89) to assess whether SBS88 could be attributed to these early phylogenetic branches. Early phylogenetic branches were defined as those of length less than 50 mutations and ceased before 50 total mutations.

**Supplementary Figure 7:**
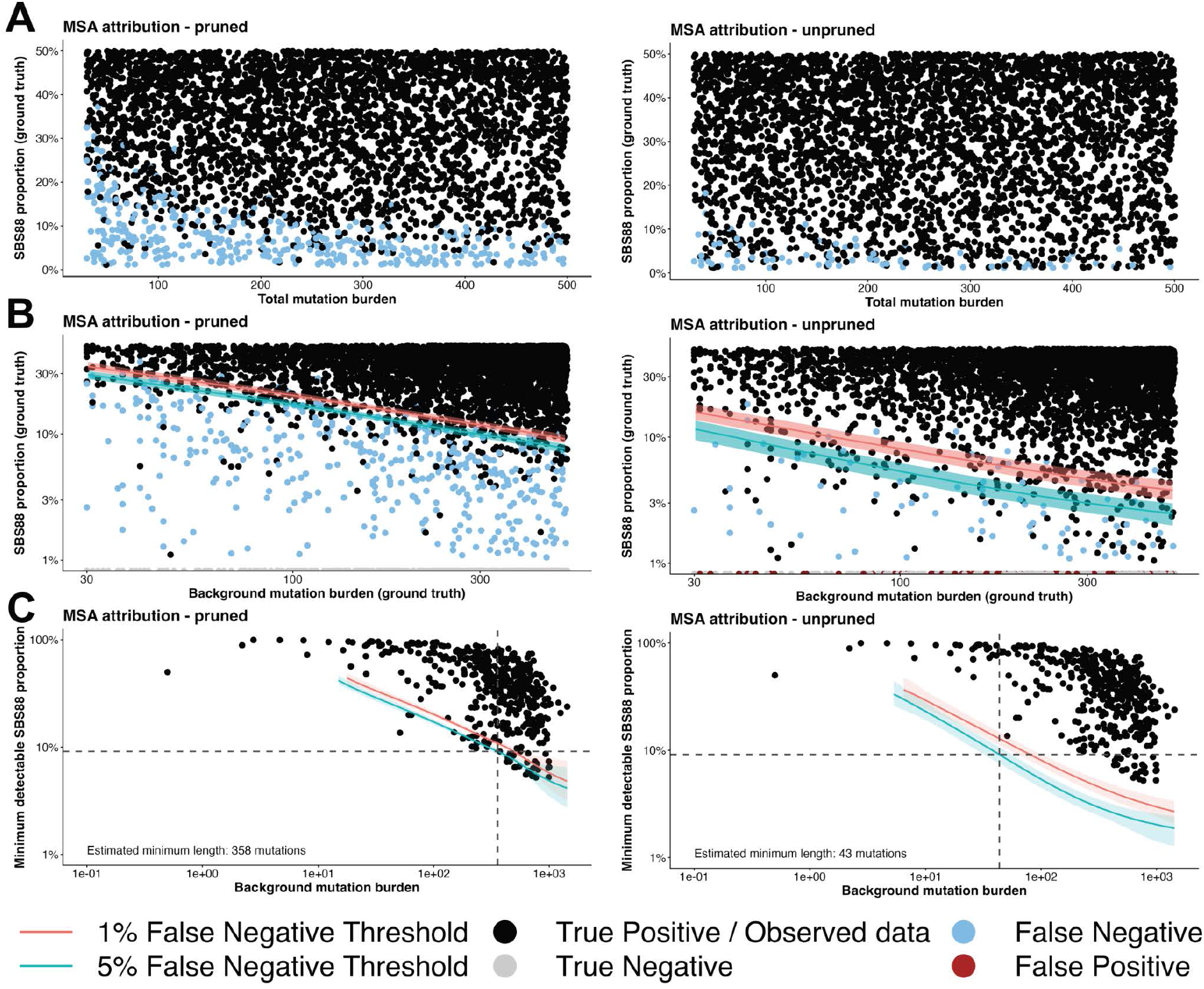
Illustration of minimum detectable branch length: **A**: Scatter plot showing attribution of SBS88 to 5500 simulated samples for pruned (left) and unpruned (right) bootstrapped attribution from MSA. Black dots indicate true positives, light blue points indicate false negatives, red points indicate false positives. **B:** Model outputs from estimation of 1% false negative threshold and 5% false negative threshold for a given background mutation burden and SBS88 proportion of mutations for pruned and unpruned outputs from MSA. 1% false negative thresholds indicate that any point above these lines have a less than 1% likelihood of being called a false negative. For a given combination of SBS88 proportion and total background mutation burden, the threshold is lower in unpruned data owing to greater sensitivity. **C:** Estimation of minimum branch length for calling a branch SBS88 negative. Dots indicate background mutation burden (x-axis) and proportion of SBS88 (y-axis) in observed branches amongst children’s crypts. The red line demonstrates 1% false negative and the blue line demonstrates 5% false negative thresholds. The minimum duration was estimated as the intersection of the lower fifth percentile of SBS88 proportion from observed data with the SBS88 negative branch < 44 mutations was deemed insufficiently long.

**Supplementary Figure 8:**
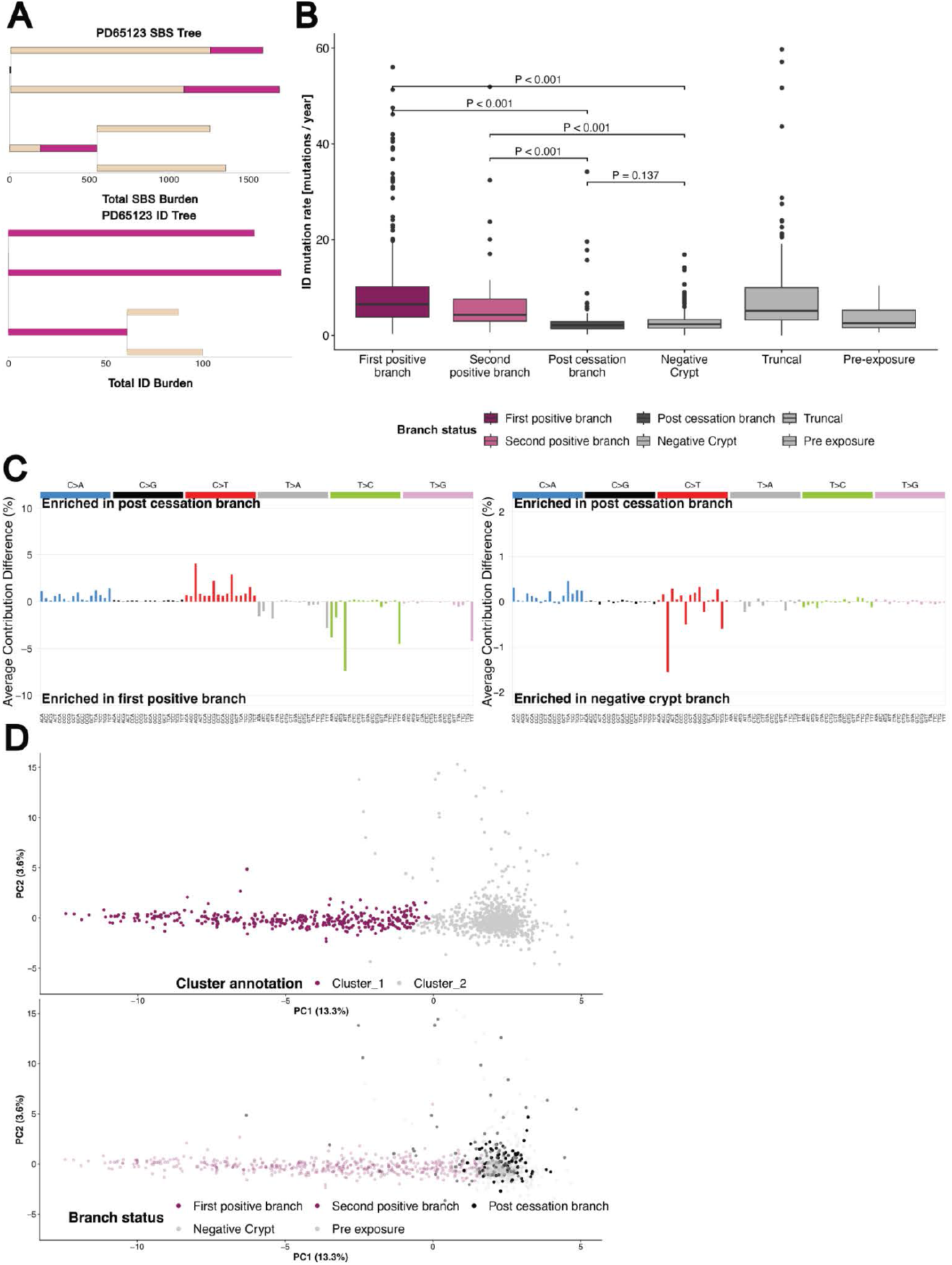
Validation of cessation amongst phylogenetic branches: **A:** Example phylogenetic tree for SBSs (top) and ID (bottom) mutations. In the SBS phylogenetic tree, the proportion of SBS mutations in each branch attributed as SBS88 annotated by pink bar, whereas the proportion of background mutations indicated by light brown. In the ID phylogenetic tree, branches are annotated as pink if SBS88 was attributed to that branch, or light brown if no SBS88 was attributed (i.e. only SBS background is attributed). **B:** Box and whisker plot of ID mutations per year with annotation of branches as either first positive, second positive, post-cessation, branches from negative crypts, truncal or pre-exposure. P-values derived from Wilcoxon rank sum test with Bonferroni correction. ID mutations per year for post-cessation branches was less than positive branches (p-value<0.001) and equivalent to negative crypts (p-value=0.137). **C:** Mutational spectra comparison for post-cessation branches to first positive branch (left) and for post-cessation branches to negative crypt branches (right). The Y-axis depicts the relative difference in proportion of trinucleotide between two branch types. Positive values indicate enrichment in post-cessation branches and negative values indicate enrichment in negative crypt branches. No trinucleotide contexts were significantly different between post cessation branches and branches from negative crypts (all pairwise T-tests p-values>0.05). **D:** Principal components analysis of mutation spectra of all branches in study analysis. Top depicts unsupervised clustering (Ward.D). Bottom shows colour annotation by branch status. Unsupervised clustering correlated with SBS88 branch annotation. Cluster 1 corresponded to SBS88 positive branches, whereas Cluster 2 was composed of SBS88 negative branches. All post-cessation branches were assigned to Cluster 2.

**Supplementary Figure 9:**
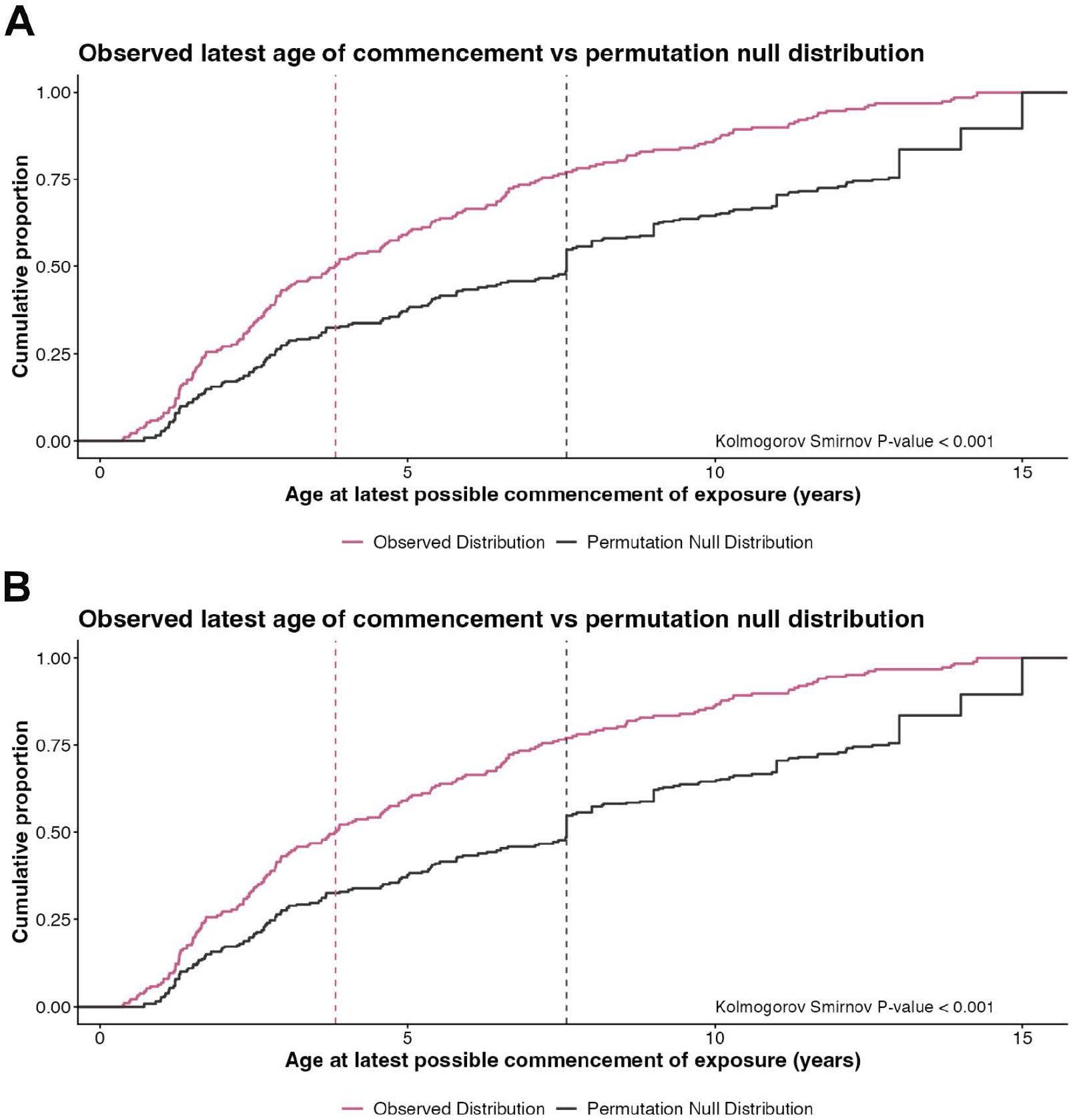
Permutation-based analyses of estimated timepoints. **A/B:** Comparison of distribution of cumulative proportion (y-axis) for age of earliest commencement of exposure (a), and age for latest possible commencement of exposure (b) between the observed distribution of timepoints (pink line) and permuted null distribution of timepoints (grey line). Permuted distributions derived from randomly permuted phylogenetic branches as SBS88 positive.

**Supplementary Figure 10:**
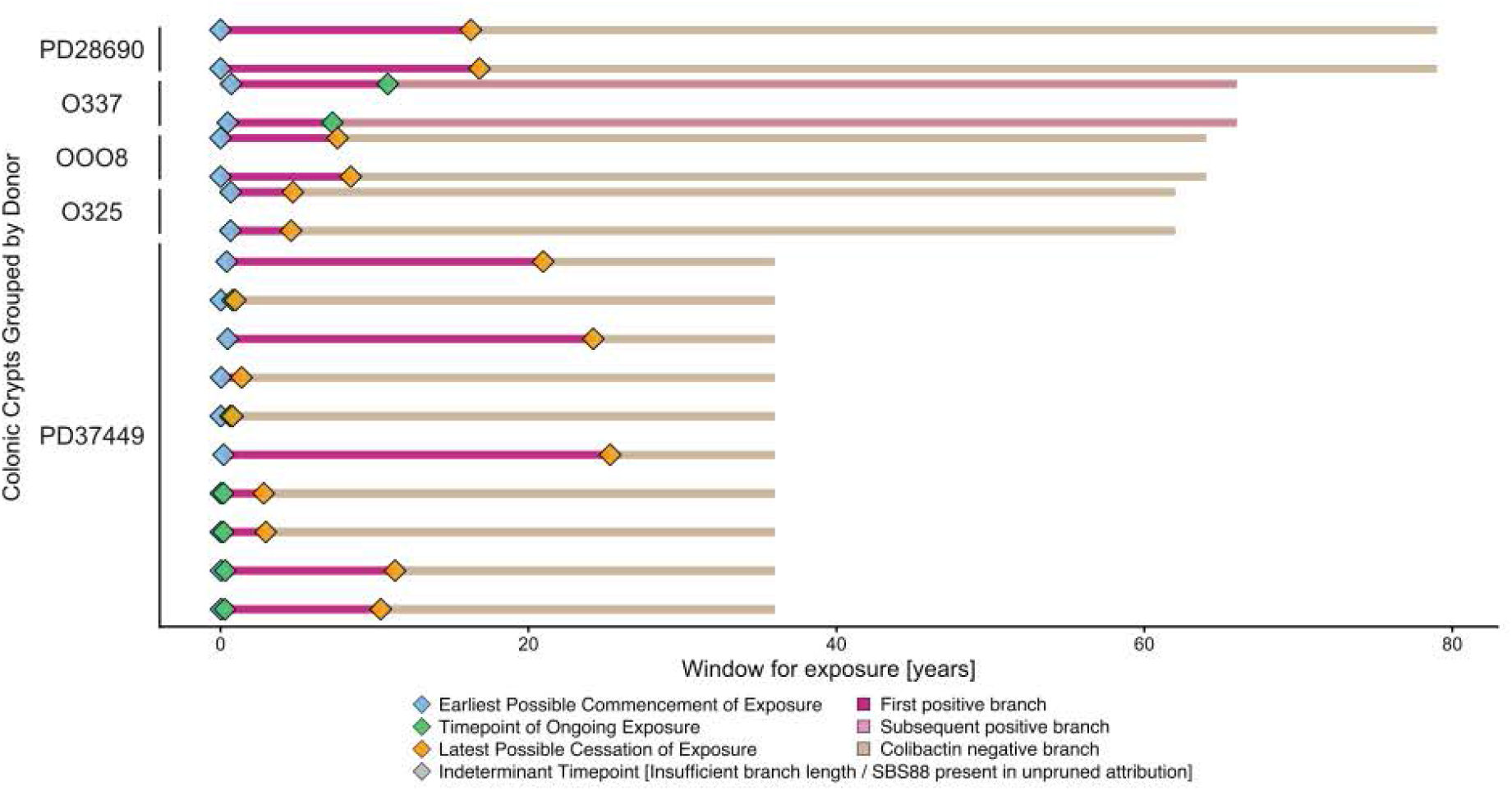
Timing colibactin mutagenesis in adult crypts. Phylogenetic estimates for window of exposure to SBS88 from n=5 individuals with informative timepoints. Blue diamonds indicate age at earliest possible commencement of exposure, green diamonds indicate ages at possible continuation of exposure, yellow diamonds indicate age at latest possible cessation of exposure. Dark pink indicates first branch per crypt positive for SBS88, light pink indicates second positive branch per crypt for colibactin mutagenesis. Light brown indicates SBS88 negative branches. All donors depicting timepoints of cessation demonstrated cessation by the age of 10.

**Supplementary Figure 11:**
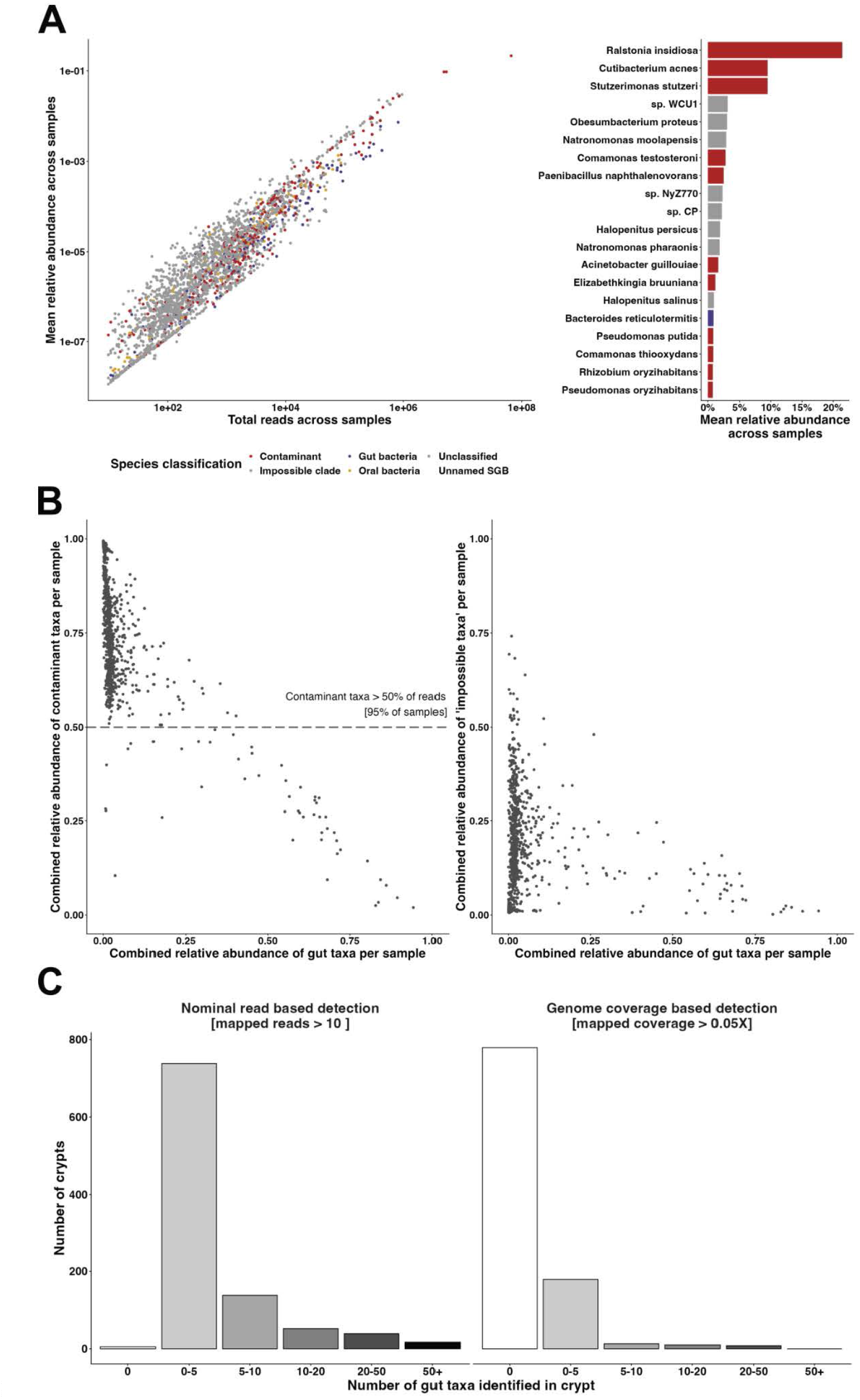
Metagenomic analysis of non-human reads. Non-human reads from individual crypts were initially mapped using krakenUniq and Metaphlan. Bacterial taxa were annotated as likely contaminants, gut taxa, or unknown. Contaminants were taxa that had previously been documented as reagent, laboratory or environmental contaminants^40,41^. **A:** High levels of contamination were observed across the dataset analysed. Left panel depicts mean relative abundance for individual species annotated by class as contaminant, gut taxa or unknown. Right panel shows the mean relative abundance for the top 20 bacterial taxa. The three most abundant taxa were all contaminants (combined assignment: 40.1% of reads). **B:** Analysis of relative abundance (y-axis) of contaminants (left panel) and biologically impossible (right panel) contaminants versus relative abundance of gut taxa per sample (x-axis). The relative abundance of contaminant-associated reads exceeded 0.5 in 95% of samples, and all samples displayed biologically impossible samples. **C:** Analysis of coverage of gut taxa per sample. Bar plot showing the number of crypts (y-axis) falling into each bin of gut taxa detected. Panels are faceted by method for taxon detection: nominal read based detection (left) and coverage based detection (right). (x-axis).

## Notes

### Competing Interest Statement

The authors have declared no competing interest.

